# Resurrecting Full-length Ancestral Schizorhodopsins and Heliorhodopsins with Structure-guided, Indel-aware Sequence Reconstruction

**DOI:** 10.64898/2025.12.28.696783

**Authors:** Haruto Ishikawa, Yasuhisa Mizutani

## Abstract

Microbial rhodopsins exhibit diverse functions ranging from ion pumps and channels to light sensors, despite sharing a common seven-transmembrane (7TM) architecture. Understanding how this functional diversity evolved is a long-standing problem, and ancestral sequence reconstruction (ASR) offers a direct route to inferring and experimentally testing plausible ancestral rhodopsins. However, ASR of 7TM proteins is often limited by alignment ambiguity and insertion–deletion (indel) uncertainty, especially in extra-membrane (EM) loops and termini. As a result, many studies focus on trimmed transmembrane (TM) cores and treat EM regions by manual curation, leaving the evolutionary history of full-length architecture difficult to test experimentally. Here we reconstruct and resurrect full-length ancestral schizorhodopsins (Anc-SzR) and heliorhodopsins (Anc-HeR), two microbial rhodopsin families that share a retinal-binding 7TM core but differ in membrane topology and EM secondary-structure elements. Starting from untrimmed alignments, we combine structure-consistent multiple sequence alignments and profile-based evolutionary models with an explicit indel-aware refinement that merges amino-acid ancestral states with binary ancestral gap inference on a fixed topology. Indel-aware refinement prevents artificially overextended ancestors and yields compact full-length sequences. AlphaFold3 predictions for the indel-corrected ancestors support high-confidence 7TM folds and recover lineage-specific EM features, including characteristic *β*-strands and short helices. Both Anc-SzR and Anc-HeR can be expressed in *Escherichia coli* and recovered as stable, colored, retinal-binding holoproteins. In a whole-cell pH assay, Anc-SzR shows light-driven proton-transport activity, whereas Anc-HeR shows no detectable ion-pumping signal, consistent with extant heliorhodopsins. Together, these results show that full-length, indel-aware ASR can produce experimentally tractable ancestral microbial rhodopsins and enable direct tests of how EM architecture evolves alongside the 7TM core.

## Introduction

Microbial rhodopsins are a broadly distributed family of retinal-binding photoreceptors found across bacteria, archaea, and microbial eukaryotes, and have also been identified in some viral genomes. Despite wide sequence divergence, they share a conserved seven-transmembrane (7TM) architecture that houses the retinal-binding pocket and the Schiff-base lysine, supporting photobiological functions such as light-driven ion transport and sensory signaling (*1*, *2*). Over the past decade, large-scale genome and metagenome sequencing has greatly expanded the catalog of microbial rhodopsins, revealing extensive functional diversity that includes proton, sodium, chloride, and other ion pumps, light-gated channels, and sensory rhodopsins, often with lineage-specific adaptations in both the transmembrane (TM) core and extra-membrane (EM) segments. This expansion raises a fundamental evolutionary question: how did such diverse photochemical and physiological functions emerge from a shared 7TM scaffold? Addressing this question requires approaches that can move beyond comparisons among extant proteins and explicitly reconstruct historical intermediates. Ancestral sequence reconstruction (ASR) offers a practical route to connecting phylogenetic history with changes in structure, spectral tuning, oligomerization, and function over evolutionary time.

ASR infers the sequences of extinct proteins at internal nodes of a phylogeny and enables experimental tests of how protein structure, stability, and function evolved over deep time (*3*, *4*). In many systems, resurrected ancestors provide tractable reagents for connecting evolutionary hypotheses to biochemical mechanisms. At the same time, ASR outcomes can be strongly affected by practical choices—most notably multiple sequence alignment (*5*, *6*), evolutionary model specification (*7*, *8*), and the treatment of insertions and deletions (indels) (*9*, *10*). Such choices can alter both tree topology and posterior probabilities (PPs) at internal nodes, yet their consequences are not always apparent from standard ASR diagnostics, motivating explicit evaluations of robustness when ancestors are taken forward for experimental resurrection.

For soluble proteins, the effects of alignment uncertainty, model choice, and indel treatment on ASR have been examined in detail (*4*, *6*). In contrast, 7TM membrane proteins present a distinct set of practical challenges that remain less systematically explored. 7TM proteins combine a conserved hydrophobic helical core with EM regions that are highly variable in both length and secondary structure, including N- and C-terminal tails, peri-membrane helix termini, amphipathic elements, *β*-strands, and flexible coils (*11*, *12*). Because these EM segments are often gap-rich and alignment-sensitive, many studies focus on trimmed TM cores and handle EM regions and termini through manual curation, thereby treating indels only implicitly (*13* –*15*). Consequently, low overall sequence identity, compositional bias, and topological constraints complicate both alignment and model fitting in many 7TM families (*16*, *17*). Together, these factors have limited efforts to reconstruct and experimentally test full-length 7TM ancestors and have left EM architecture underused as an explicit source of evolutionary information.

As demonstrated in previous studies, alignment errors can propagate into ASR errors and distort down-stream structural and functional inferences, sometimes more strongly than phylogenetic uncertainty itself (*18*, *19*). In particular, Aadland and Kolaczkowski identified insertion–deletion (indel) errors as a major contributor to ASR inaccuracy across realistic simulation conditions and emphasized that many practical workflows still handle indels only implicitly through the choice of a single alignment rather than by explicit modeling (*19*). This concern is amplified in 7TM proteins, where EM regions are often long and gap-rich: such datasets provide a stringent setting for evaluating how alignment and indel uncertainty affect full-length ancestral reconstructions that are intended for experimental testing.

Schizorhodopsins (SzRs) and heliorhodopsins (HeRs) share a conserved 7TM architecture and the retinal-binding lysine in the seventh transmembrane helix (TM7), yet differ sharply in membrane topology and oligomeric state: SzRs adopt an N-out/C-in orientation and form trimers, whereas HeRs display an inverted N-in/C-out topology and form dimers (*20* –*25*). Functionally, SzRs include inward proton pumps, whereas HeRs are typically considered non-pumping photoreceptors, providing a sharp functional contrast in addition to topology and oligomeric state (*2*, *20*, *22*). Both families also contain characteristic EM secondary-structure elements—including short helices and short or long *β*-strands—that contribute to intersubunit packing and function (*21*, *23* –*25*). These contrasts provide a controlled setting to (i) compare alignment and model choices, (ii) quantify the impact of indels on full-length reconstructions, and (iii) test whether ancestral sequences can recover not only the 7TM core but also lineage-specific EM architectural motifs with high confidence. (Here and below, TM1–TM7 denote the seven canonical transmembrane helices numbered from the N terminus.)

Recent advances now make it possible to revisit these questions for SzRs and HeRs more systematically. First, profile-based substitution models such as Q.pfam+R better reflect the compositional and site-specific constraints typical of membrane proteins than classical empirical matrices (*16*), and are now readily usable in routine likelihood-based analyses. Second, explicit binary gap models implemented in RAxML and related tools provide a principled route to reconstructing ancestral indel patterns on a fixed topology (*26*, *27*). Third, structure prediction with residue-wise confidence measures (pLDDT), such as those provided by AlphaFold (*28*), offers an independent way to evaluate whether inferred ancestral sequences yield coherent 7TM folds and plausible extra-membrane architectures. Together, these developments enable full-length ancestral reconstructions to be assessed not only by evolutionary fit but also by structural plausibility—an essential step when ancestors are designated for experimental resurrection.

Accordingly, we reconstruct and experimentally resurrect full-length ancestral schizorhodopsins (Anc-SzR) and heliorhodopsins (Anc-HeR) and evaluate the reliability of both TM and EM regions in this study. Starting from untrimmed, structure-consistent alignments, we infer amino-acid ancestral states under profile-based models, refine ancestral sequences with explicit indel-aware masking based on binary gap reconstructions on a fixed topology, and assess structural plausibility using AlphaFold predictions for monomeric and oligomeric assemblies. To summarize node-level robustness in a transparent way, we report joint sequence–structure confidence by mapping PPs together with pLDDT and interpret these alongside bootstrap support for the corresponding splits.

We heterologously express and purify indel-corrected Anc-SzR and Anc-HeR in *Escherichia coli* (*E. coli*), yield stable retinal-binding holoproteins with rhodopsin-like spectra, consistent with characteristic visible absorption spectra. Finally, we provide initial functional readouts using a whole-cell pH assay: Anc-SzR shows a protonophore carbonyl cyanide m-chlorophenyl hydrazone (CCCP) sensitive light-driven pH response, whereas Anc-HeR shows no detectable pumping signal, consistent with extant HeRs. Using these resurrected ancestors as a readout, we evaluate how alignment choice, substitution models, and explicit indel-aware refinement influence node stability, sequence length, and confidence in EM regions, and we compare evolutionary support with structure-based confidence across ancestral depths. Together, our results establish SzR and HeR as a tractable system for full-length 7TM resurrection and show that EM architecture can be reconstructed and evaluated as an explicit component of membrane-protein evolution rather than being routinely trimmed away.

## Materials and Methods

### Sequence datasets and curation

HeR and SzR sequences were compiled by combining previously published datasets with additional database searches. Previously characterized HeR and SzR sequences were taken from Inoue et al. and Bulzu et al. (*22*, *29*) and supplemented with homologs identified by BLAST searches against NCBI protein databases and by profile-based searches of MGnify metagenome assemblies (*30*). Because SzR-like sequences are sparsely represented in standard protein databases, most additional SzR candidates were obtained from metagenomic assemblies. Several HeR sequences were directly taken from the metagenome-derived dataset of Bulzu et al. (*29*); for these entries, we retained the original contig/gene identifiers when a unique public accession could not be assigned. Although light-driven proton pumping has been reported for some giant-virus heliorhodopsins (*31*), we did not intentionally include those previously reported viral heliorhodopsin sequences in our curated dataset. Because metagenome assemblies can, in principle, contain unannotated viral contigs, we cannot exclude an unrecognized viral origin for a minority of metagenome-derived entries; however, no sequences retained after curation were annotated as viral in their source records. All amino-acid sequences used in this study are provided as Supplementary FASTA files.

Candidate sequences were filtered in four stages: (i) *Length filtering*: sequences shorter than 200 amino acids for HeRs or 150 amino acids for SzRs were discarded to remove obvious fragments and mispredicted multi-pass membrane proteins. (ii) *Functional-residue filtering*: sequences that did not contain any lysine residues were excluded. (iii) *Redundancy reduction*: highly similar sequences (>95% pairwise identity) were clustered with CD-HIT (*32*), retaining one representative per cluster. (iv) *Phylogenetic diversity filtering*: Treemmer (*33*) was applied to a preliminary tree to downsample near-identical sequences while preserving long internal branches.

After automated filtering, we performed a topology-consistency screen using the PSI/TM-Coffee transmembrane-mode alignment (*34*). Because microbial rhodopsins have a conserved 7TM core, we required that the alignment contained continuous, gap-poor registers across TM1–TM7. Sequences were excluded if they showed obvious loss of TM registers (e.g., long internal deletions or extensive gaps within one or more predicted TM helices), or if conserved core positions—including the Schiff-base lysine in TM7—were missing due to truncation or misprediction. This curation step was applied only to remove sequences incompatible with a canonical 7TM rhodopsin fold; no additional per-sequence TM-helix prediction beyond the alignment-based TM-mode constraints was performed. The final non-redundant dataset contained 228 sequences (115 HeRs, 110 SzRs and 3 outgroup microbial rhodopsins). Two reduced datasets, HeR+outgroup and SzR+outgroup, were derived from this set, sharing the same outgroup panel. Sequence lengths ranged from approximately 195 to 305 residues, consistent with known 7TM microbial rhodopsin architectures.

### Multiple sequence alignment

Multiple sequence alignments were generated for the full HeR+SzR+outgroup dataset and for the two subset datasets (HeR+outgroup and SzR+outgroup) using PSI/TM-Coffee and MAFFT (*34*, *35*). Unless otherwise stated, default parameters of the respective programs or web servers were used.

Structure-aware alignments were obtained with PSI/TM-Coffee in transmembrane mode using UniRef100 as the homologous sequence database (*34*). In this mode, predicted transmembrane segments and additional homologous sequences are incorporated in the consistency-based alignment procedure, stabilizing TM/EM boundaries. The resulting alignments were exported in full-length FASTA format without manual editing of internal regions.

As a widely used non–structure-guided baseline, we generated MAFFT alignments using the L-INS-i strategy (*35*). For exploratory analyses and for a classical workflow comparison, additional alignments were generated with the faster FFT-NS-1 strategy in MAFFT; because FFT-NS-1 showed visible mismatches at TM/EM boundaries and yielded unstable HeR reconstructions in some conditions, it was used only in sensitivity analyses and in the classical pipeline comparison (see below).

To examine the influence of mild gap-based trimming, we also produced trimmed versions of each alignment using trimAl with the gappyout option (*36*). These trimmed alignments were used primarily to assess robustness of model selection and topology. In addition, to provide a conventional column-trimming baseline for the impact of indel-aware refinement, we performed IQ-TREE ASR on the gappyout-trimmed alignments and carried these gap-unaware ancestors forward to AlphaFold evaluation in the same manner as the untrimmed ASR results. All primary indel-aware reconstructions and downstream analyses otherwise relied on untrimmed, full-length alignments (i.e. no aggressive manual trimming of loops or termini was applied).

### Phylogenetic inference and model selection

Maximum-likelihood (ML) phylogenies were inferred with IQ-TREE v3.0.1 (*37*) for the HeR+SzR+outgroup dataset and for the HeR+outgroup and SzR+outgroup subsets. Alignments were analyzed as single-partition protein datasets.

For each alignment, ModelFinder (*38*) (-m MFP) was used to compare empirical matrices (e.g. LG, WAG, JTT) with various rate-heterogeneity schemes and profile families including Q.pfam and related models.

Model fit was assessed using the Bayesian information criterion (BIC), and the model with the lowest BIC was selected. In the main analyses we adopted PSI/TM-Coffee with the best-fit Q.pfam+R model (typically Q.pfam+R7) as the baseline evolutionary framework (see Results and Table 1), and used MAFFT L-INS-i as an alignment sensitivity check.

**Table 1:** Summary of datasets, alignments, and best-fit substitution models used for phylogenetic inference and ancestral sequence reconstruction.

| Dataset | Alignment strategy | $N$ sequences | Alignment length<br>(residues) | Best-fit model<br>(BIC) | Notes |
| --- | --- | --- | --- | --- | --- |
| HeR+SzR+outgroup | PSI/TM-Coffee | 228 | 449 | Q.pfam+R7 | Baseline dataset used for joint HeR–SzR phylogeny and ASR. Outgroup sequences were constrained as the external clade. |
| HeR+SzR+outgroup | MAFFT L-INS-i | 228 | 421 | Q.pfam+R7 | Alternative alignment used to assess alignment sensitivity under the same model family. |
| HeR+outgroup | PSI/TM-Coffee | 118 | 533 | Q.pfam+R6 | HeR-only subtree used for node-level analyses of Anc-HeR. |
| HeR+outgroup | MAFFT L-INS-i | 118 | 401 | LG+F+R6 | Alignment/model combination that yielded a near-polytomy around the HeR ancestor in UFBoot2 consensus trees. |
| SzR+outgroup | PSI/TM-Coffee | 113 | 320 | Q.pfam+R5 | SzR-only subtree; Anc-SzR showed high and stable bootstrap support. |
| SzR+outgroup | MAFFT L-INS-i | 113 | 287 | Q.pfam+R6 | Alignment used to confirm robustness of Anc-SzR to alignment strategy. |

Best-fit models are those selected by ModelFinder under the Bayesian information criterion (BIC).

ML tree searches were run in IQ-TREE with ultrafast bootstrap approximation (1,000 replicates) and SH-aLRT support (1,000 replicates) (*39*). Three outgroup rhodopsins were specified to root the tree. The best-scoring ML tree (.treefile) was used for visualization and as the fixed topology in downstream ASR and binary indel reconstructions. The UFBoot2 consensus tree (.contree) was used to determine whether focal branches were resolved or collapsed and to obtain the branch support values used in node-level summaries. UFBoot2 support values used in the main text and Table 2 were taken from the consensus trees (.contree) generated under the baseline settings. To assess convergence, we repeated UFBoot2 analyses with increasing replicate numbers and an independent run using an alternative random seed (Table S4).

**Table 2:** Composite node-level reliability metrics integrating posterior probabilities (PP), AlphaFold confidence (pLDDT), and ultrafast bootstrap support (UFBoot) for key ancestral nodes.

| Alignment | Ancestor | UFBoot2 (ML tree)<br>(%) | UFBoot2 (consensus)<br>(%) | Mean PP<br>(%) | Mean pLDDT | Mean (PP×pLDDT) | Composite score<br>(0–100) |
| --- | --- | --- | --- | --- | --- | --- | --- |
| PSI/TM-Coffee | Anc-SzR | 99 | 99 | 81.6 | 95.5 | 78.1 | 77.3 |
|  | Anc-HeR | 100 | 54 | 90.6 | 95.9 | 87.2 | 47.1 |
|  | Anc-SH | 100 | 100 | 54.9 | 93.2 | 51.6 | 51.6 |
| MAFFT L-INS-i | Anc-SzR | 96 | 96 | 80.4 | 94.3 | 76.1 | 73.0 |
|  | Anc-HeR | 100 | 0<sup>a</sup> | 91.4 | 93.6 | 86.0 | 0.0<sup>a</sup> |
|  | Anc-SH | 100 | 100 | 57.5 | 93.6 | 54.3 | 54.3 |

UFBoot2 (ML tree) values are those reported on the best-scoring ML topology (**.treefile**), whereas UFBoot2 (consensus) are frequencies on the bootstrap consensus tree (**.contree**). Mean PP, mean pLDDT, and mean PP×pLDDT are reported on a 0–100 scale. Composite scores were computed as $100 \times b \times \overline{PP \times pLDDT}$ , where $b$ is the consensus UFBoot2 support expressed as a fraction ( $b = \text{UFBoot2}/100$ ) and $\overline{PP \times pLDDT}$ is the mean product on a 0–1 scale.
<sup>a</sup> For MAFFT L-INS-i, the HeR ancestral branch collapses in the UFBoot2 consensus tree (polytomy), so UFBoot2 support is effectively 0 and the composite score is 0 by definition for this alignment.

To benchmark the impact of increased model complexity, additional trees were inferred under LG+C60+F+R8 using otherwise identical settings. These analyses were treated as robustness checks rather than the primary framework for ASR.

### Ancestral sequence reconstruction in IQ-TREE

ASR was performed in IQ-TREE v3.0.1 on the ML topologies inferred under the baseline Q.pfam+R model. For each dataset (HeR+SzR+outgroup, HeR+outgroup, SzR+outgroup) and each alignment (PSI/TM-Coffee, MAFFT L-INS-i), we carried out marginal ASR at all internal nodes using –ancestral, treating the topology as fixed.

IQ-TREE reports, for each internal node and alignment column, posterior probabilities (PPs) over the 20 amino-acid states and the maximum-posterior state. These values were extracted from the .state files.

We focused on three internal nodes in the full tree: the SzR ancestor (Anc-SzR), the HeR ancestor (Anc-HeR), and their MRCA (Anc-SH). Sensitivity analyses under LG+C60+F+R8 were performed on the same alignments and topologies to assess model-dependent variation in ancestral states and PP distributions (see Results).

Unless otherwise noted, PP-based summaries and all structural evaluations reported in this study refer to the indel-corrected ancestral sequences obtained after the indel-aware refinement described below.

### Indel-aware reconstruction using binary gap models

To make indel treatment explicit and node specific, we implemented a two-stage refinement pipeline inspired by binary gap coding approaches (*27*). In this workflow, IQ-TREE is used for amino-acid ASR, and RAxML-HPC v8.2.12 (*26*) is used for binary gap inference on the same fixed topology.

*Binary gap inference.* For each amino-acid alignment, we generated a binary gap matrix by recoding each column as 1 (residue present) or 0 (gap). RAxML-HPC was then used to infer ancestral gap states under a two-state (gap/non-gap) model on the fixed IQ-TREE ML topology. This yields, for each internal node and alignment column, a binary ancestral call for presence/absence.

*Node mapping and node-specific masking.* Because IQ-TREE and RAxML assign different internal node labels, we mapped nodes by bipartitions: for each internal node, the set of descendant tips defines a split, and we matched IQ-TREE nodes to RAxML nodes by exact agreement of these leaf-set bipartitions (with outgroup-aware handling of complementary splits). For each ancestor and alignment column, if the binary reconstruction indicated a gap state (0), the corresponding IQ-TREE amino-acid state and PP were masked and removed; if a residue state (1) was inferred, the IQ-TREE maximum-posterior residue and its PP were retained. Indel-corrected ancestral sequences were then assembled by concatenating unmasked positions in their original alignment order.

All downstream analyses (sequence identity comparisons, regional PP/pLDDT summaries, composite scores, and structure prediction) used these indel-corrected sequences. Gap recoding, RAxML parsing, bipartition-based node mapping, and masking were implemented in custom Python scripts (see Software and reproducibility).

### Extant sequence reconstruction proxy validation

To validate reconstruction accuracy on sequences of known ground truth, we performed an extant-sequence reconstruction (ESR) proxy test on SzR4 and HeR 48C12 under the baseline setting (PSI/TM-Coffee; Q.pfam+R7) (*40*). For each target tip, we replaced the original leaf by a proxy subtree with two dummy leaves (suffixes _A and _B) attached by long branches, while keeping the internal proxy node label identical to the original target. The initial dummy-branch length was set to 500. Because fixing branch lengths is not supported under FreeRate site-rate models in IQ-TREE (e.g., Q.pfam+R), we constrained the topology (-te) but allowed branch lengths to be re-optimized by maximum likelihood during the ESR runs. We selected the initial dummy branch length of 500 after confirming that ESR identity scores plateaued at large dummy branch lengths (tested 10–2000), consistent with saturation of proxy-leaf decoupling. IQ-TREE ASR was then run on the fixed proxy topology (with branch lengths optimized), and the reconstructed state at the proxy internal node was compared to the true target sequence at sites where the true residue was not a gap. We report (i) identity between the maximum-posterior state and the true residue and (ii) the mean posterior probability assigned to the true residue, globally and by TM/EM regions.

### AlphaFold structure prediction and confidence metrics

Predicted structures for ancestral and extant rhodopsins were obtained using an AlphaFold3 web server (*28*). For single-chain modeling, each sequence was submitted as a single input chain; for oligomeric modeling, identical chains were provided as multiple sequences in the same submission to obtain homomeric complexes (stoichiometries of 1, 2, 3, and 5). In this web workflow, multimeric prediction is therefore specified by multi-sequence input rather than by an explicit “Multimer” option.

For each submission, five models were generated, and the top-ranked model (server ranking) was used for downstream analyses. Per-residue pLDDT values were extracted from the B-factor field of the output coordinate files; analyses were restricted to C*_α_* atoms, yielding one pLDDT value per residue. These pLDDT values were paired with site-wise PPs from IQ-TREE using alignment coordinates, restricting the mapping to residues retained after indel-aware masking.

To assess oligomeric preferences, Anc-SzR, Anc-HeR, and Anc-SH were modeled as homomeric assemblies with stoichiometries of 1 (monomer control), 2 (dimer), 3 (trimer), and 5 (pentamer). We focused on 2/3/5 because microbial rhodopsins are experimentally observed as dimers, trimers, or pentamers, whereas tetramers have not been reported. For each stoichiometry and ancestor, we recorded pTM and ipTM for the top-ranked model, which quantify predicted fold accuracy and interface quality. Structural inspection and figure preparation were performed in UCSF ChimeraX and PyMOL.

### Definition of TM and EM regions using OPM

To quantify region-specific reliability, we partitioned each ancestral model into transmembrane (TM) and extra-membrane (EM) regions using membrane embeddings from the Orientations of Proteins in Membranes (OPM) database (*41*). For SzR, we used the OPM entry for SzR4 (PDB 7e4g). For HeR, we used the OPM entry corresponding to the HeR 48C12 structure (PDB 6uh3) used for TM/EM boundary definition. Residues lying within the hydrophobic core of the OPM membrane slab were classified as TM, and all remaining residues were classified as EM. In this convention, EM includes N- and C-terminal tails, perimembrane helix termini, *β*-strands, short coils, and amphipathic helices outside the membrane core.

TM/EM labels were transferred to reconstructed ancestors via the same structure-consistent alignments used in ASR. Labels were propagated column-wise from the extant template sequence to the ancestral sequence; sites aligned to gaps in the ancestor or removed by indel-aware masking were excluded. Using these labels, we computed region-level means for PP, pLDDT, PP×pLDDT, and PP−pLDDT for TM and EM residues and, where indicated, for individual TM helices and adjacent EM segments.

### Composite reliability metric

To integrate evolutionary, statistical, and structural information at the node level, we defined a composite reliability metric for Anc-SzR, Anc-HeR, and Anc-SH. For each ancestor, site-wise marginal PPs were extracted from IQ-TREE .state files and mapped onto the indel-corrected sequence; pLDDT values were taken from the corresponding AlphaFold 3 model and normalized to 0–1.

Let PP*_i_* and pLDDT*_i_* denote the posterior probability and normalized pLDDT (0–1) for residue *i*, and let *N* be the number of C*_α_* sites considered. We defined

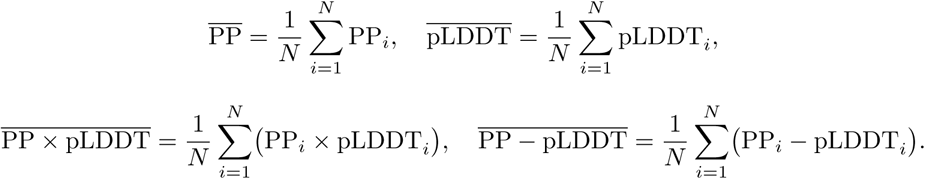

In the Results, these quantities are reported on a 0–100 scale by multiplying by 100.

Bootstrap support (BS) for each ancestor was taken from the UFBoot2 consensus tree (.contree) under the corresponding alignment and model. BS was defined as the fraction of bootstrap trees containing the relevant split, expressed as *b* = BS*/*100 on a 0–1 scale. When a branch was collapsed in the consensus (i.e. the split was absent), *b* was set to 0.

The composite node-level score was defined as

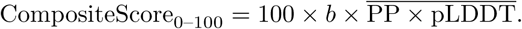

Composite scores were computed separately for each alignment/model combination; baseline values correspond to PSI/TM-Coffee with Q.pfam+R7.

### Expression, purification and spectroscopy of ancestral rhodopsins

Genes encoding Anc-SzR and Anc-HeR were synthesized as contiguous open reading frames corresponding exactly to the inferred amino-acid sequences (GenScript), without additional fusion partners or stabilizing mutations. For Anc-SzR, the gene was cloned into the NdeI and XhoI sites of pET-21a(+), placing the ASR-derived open reading frame in frame with the vector-encoded C-terminal His_6_ tag. For Anc-HeR, the gene was cloned into the NdeI and BamHI sites of pET-15b, such that the vector-encoded N-terminal His_6_ tag and thrombin recognition sequence precede the ASR-derived open reading frame and the C terminus corresponds exactly to the inferred ancestral sequence. For both constructs, the initiator methionine is encoded by the ATG within the NdeI site (CATATG) and is counted as the first residue of the ASR-derived sequence.

Recombinant Anc-SzR and Anc-HeR were expressed in *E. coli* C43(DE3). Cultures were grown in LB medium supplemented with ampicillin at 37 °C and induced at OD_600_ ≈ 1.0 with 1 mM IPTG together with 10 *µ*M all-*trans* retinal (ethanol stock). After induction, cultures were incubated for 20 h at 25 °C with shaking, which yielded strongly pigmented cell pellets for both ancestors (see Results). For qualitative comparison of expression conditions, additional cultures were induced at 37 °C for 4 or 20 h.

For purification, cells were harvested by centrifugation, resuspended in buffer, and disrupted by sonication. Membrane fractions were collected by ultracentrifugation, resuspended in detergent-containing buffer, and solubilized with n-dodecyl-*β*-D-maltoside (DDM). Solubilized pigments were purified by Co^2+^-affinity chromatography and eluted in imidazole-containing buffer with DDM. Eluted fractions were exchanged into a Tris/NaCl buffer containing DDM for spectroscopic measurements. Both Anc-SzR and Anc-HeR showed anomalously low apparent molecular weights by SDS–PAGE compared with calculated masses, despite migrating as single, well-resolved bands (Fig. S5). Detailed buffer compositions and chromatography conditions are provided in the Supplementary Methods.

UV–visible absorption spectra were recorded at room temperature using a 10 mm path length quartz cuvette. Spectra were collected over 300–700 nm with the corresponding buffer used for baseline correction.

### Whole-cell pH assay

Light-driven ion-transport activity was assessed in whole cells by monitoring external pH changes upon illumination (*42*). *E. coli* C43(DE3) expressing Anc-SzR or Anc-HeR were washed and resuspended in 300 mM NaCl (no buffering agent) and illuminated at 540 nm. Where indicated, the protonophore CCCP (30 µM) was added prior to illumination. pH traces were recorded for dark–light–dark cycles. For visualization in the main text (Fig. 8), traces were baseline-corrected by subtracting the pH value at the end of the pre-illumination dark period, and plotted as ΔpH. Raw pH traces without baseline correction are shown in Fig. S6. Additional experimental details are provided in the Supplementary Methods.

### Classical RAxML+PAML workflow for comparison

As a classical reference point, we also performed ASR using a RAxML+PAML workflow. ML trees were inferred with RAxML-NG v1.2.2 (*43*) under an empirical amino-acid model with among-site rate heterogeneity (typically LG+F+G4). These topologies were then supplied to codeml (PAML v4.10.7) (*44*) for marginal ASR under a fixed LG model with gamma-distributed rate variation (four categories). Full control files and parameter settings are provided as Supplementary Data. Because the classical workflow uses different substitution models from the baseline IQ-TREE analyses (LG-based versus Q.pfam-based profile models), differences in reconstructed ancestors under this comparison reflect a combination of software and model effects rather than a pure implementation effect.

### Software and reproducibility

Sequence curation used CD-HIT v4.8.1 (*32*) and Treemmer v0.3 (*33*). Multiple sequence alignments were generated with MAFFT (L-INS-i and FFT-NS-1) v7.525 (*35*) and PSI/TM-Coffee v11.0 in transmembrane mode (*34*). Alignment trimming used trimAl v1.5 (*36*). ML phylogenies, model selection, and ASR were performed with IQ-TREE v3.0.1 (*37*); binary gap reconstructions used RAxML-HPC v8.2.12 (*26*); classical comparisons used RAxML-NG v1.2.2 (*43*) and PAML v4.10.7 (*44*). Structural predictions were obtained using an AlphaFold3 web server (*28*), and visualization/figure preparation used UCSF ChimeraX v1.10.1 (*45*) and PyMOL v3.1.0 (Schrödinger, LLC).

Custom Python 3 scripts were written to (i) convert alignments to binary gap matrices, (ii) map RAxML ancestral gap states to IQ-TREE node labels by bipartitions, (iii) generate indel-corrected ancestral sequences, and (iv) compute PP/pLDDT-derived summaries (PP, pLDDT, PP×pLDDT, PP−pLDDT, composite scores). These scripts, configuration files, and example command lines are distributed as the Consis-tASR pipeline and as an archival snapshot (see Supporting information).

## Results

### Alignment and model choice shape the stability of Anc-HeR but not Anc-SzR

Multiple sequence alignment had a strong impact on the stability of HeR–SzR phylogenies, with the effect most pronounced in the HeR lineage. We therefore compared a structure-aware strategy (PSI/TM-Coffee) (*34*), which incorporates transmembrane-specific constraints and homology information, a widely used high-accuracy baseline (MAFFT L-INS-i) (*35*) for protein families that are difficult to align. PSI/TM-Coffee consistently preserved TM topology and EM registers across the dataset, whereas L-INS-i produced broadly coherent alignments but introduced localized ambiguity around the HeR TM1–TM2 region. As demonstrated in the following section, these localized differences translate into marked differences in topological stability around Anc-HeR, while Anc-SzR remains robust across alignment choices.

For each alignment, we used ModelFinder in IQ-TREE (*37*, *38*) to select the best-fit amino-acid substitution model under the Bayesian information criterion (BIC). Across the full HeR+SzR+outgroup dataset and the two reduced subsets (HeR+outgroup and SzR+outgroup), profile models from the Q.pfam family with FreeRate heterogeneity (Q.pfam+R5–R7) were consistently favored (Table 1). In the full dataset, Q.pfam+R7 was selected for both PSI/TM-Coffee and MAFFT L-INS-i alignments and lowered BIC by roughly 200–250 units relative to the best LG-based FreeRate alternatives, consistent with strong support for a membrane-aware profile model over conventional empirical matrices.

However, a single exception was observed in the MAFFT L-INS-i alignment of the HeR+outgroup dataset. In this case, an LG+F+R model was favored, and the HeR ancestral branch collapsed into a near-polytomy in the UFBoot2 consensus tree, indicating locally unstable support around this split (*39*). In our analyses, this behavior underscores that alignment quality can dominate model choice in difficult 7TM datasets: when the alignment introduces local inconsistencies—most notably around the HeR TM1–TM2 registers—both the preferred substitution model and the resolution of nearby internal nodes can shift. The overall preference for Q.pfam+R models was unchanged when moderately gapped terminal regions were removed with trimAl gappyout, suggesting that the signal favoring Q.pfam+R is not driven solely by gap-rich columns but is supported across both membrane-embedded and extra-membrane regions (Table S1).

For completeness, we also tested a more parameter-rich empirical profile-mixture model (LG+C60+F+R8), which improved overall fit relative to Q.pfam+R7 (Table S1). However, improved fit did not translate into improved stability of the alignment-sensitive HeR region: under LG+C60+F+R8, the Anc-HeR branch remained weak or unstable in the UFBoot2 consensus (absent for PSI/TM-Coffee and 47% for MAFFT L-INS-i), whereas Anc-SzR and Anc-SH, the most recent common ancestor (MRCA) of schizorhodopsin and heliorhodopsin, remained strongly supported. Because LG+C60+F+R8 is substantially more computationally demanding and did not change the qualitative topology or node-level reliability patterns reported below, we used Q.pfam+R7 as the primary model and treated LG+C60+F+R8 as a robustness check (Tables S2 and S3). It is important to note that the consistent preference for Q.pfam+R models also dictated the methodological choice of performing ASR in IQ-TREE, as profile models of this class are not available in classical pipelines based on RAxML or PAML. A direct comparison with a conventional RAxML+PAML workflow under LG-based models is provided in Tables S2 and S3.

These patterns are summarized in Table 1 and visualized in Fig. 1, highlighting the strong dependence of Anc-HeR on alignment quality and the robust behavior of the SzR lineage across all conditions. Because HeR topology was alignment-sensitive, we explicitly assessed UFBoot2 convergence across replicate counts and random seeds (Table S4). Accordingly, unless stated otherwise, downstream analyses used PSI/TM-Coffee (Q.pfam+R7) as the baseline.

**Figure 1:**
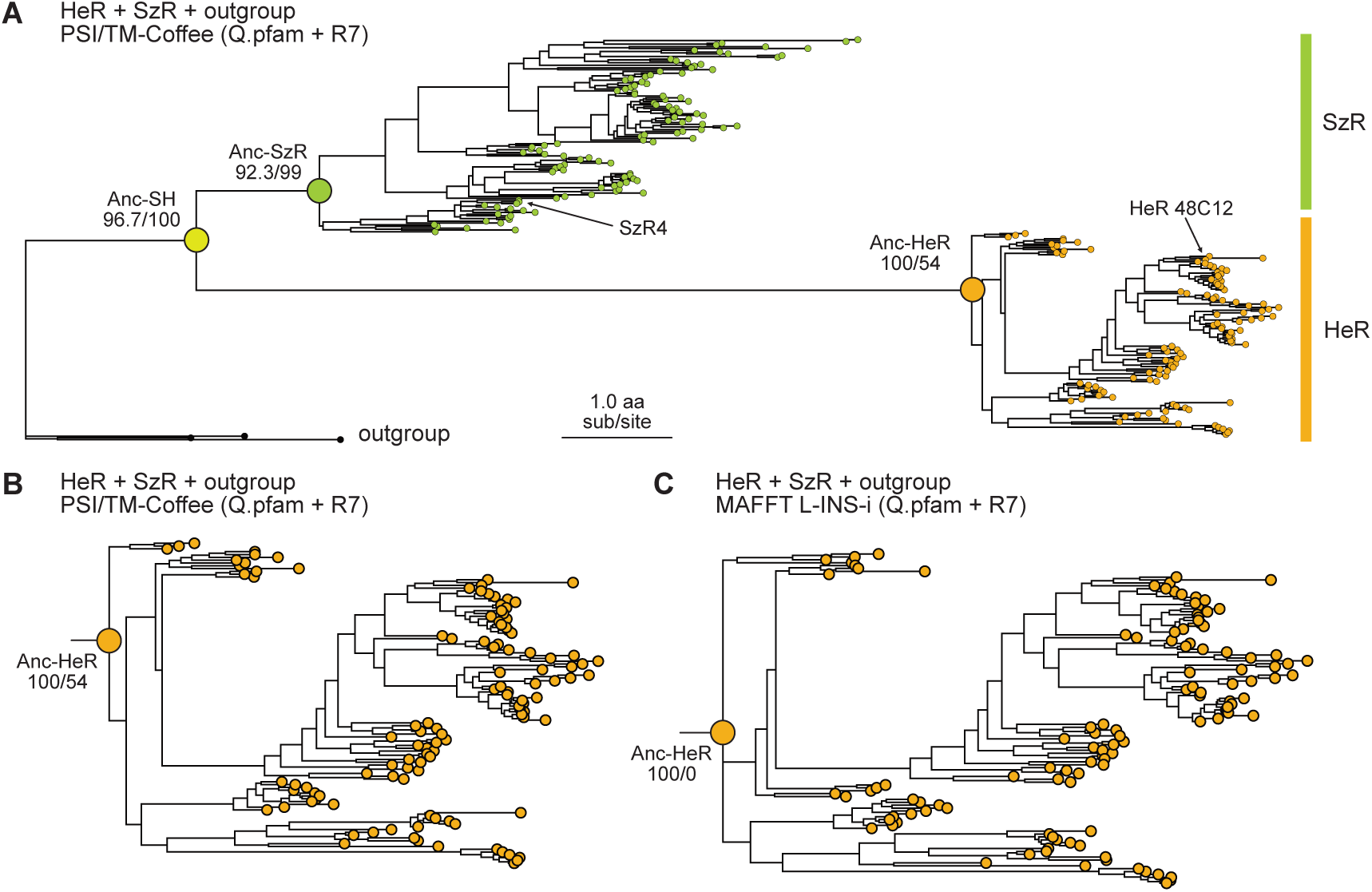
Phylogenetic context of SzR and HeR ancestors and alignment sensitivity of the HeR lineage. (A) Maximum-likelihood (ML) tree for the HeR+SzR+outgroup dataset inferred from the PSI/TM-Coffee alignment under Q.pfam+R7. HeR (orange), SzR (green), and outgroup clades are indicated, and the three focal ancestral nodes (Anc-SH, Anc-SzR, and Anc-HeR) are labeled with SH-aLRT/UFBoot2 support. Extant structural representatives (SzR4 and HeR 48C12) are marked. (B) Enlargement of the HeR clade from (A), highlighting internal branching patterns and support values around Anc-HeR under PSI/TM-Coffee. (C) Corresponding enlargement of the HeR clade inferred from the MAFFT L-INS-i alignment under the same model (Q.pfam+R7), showing reduced resolution around Anc-HeR and collapse of the corresponding split in the UFBoot2 consensus. Support values are shown as SH-aLRT/UFBoot2. When a split is absent from the UFBoot2 consensus tree (i.e., collapsed into a local polytomy), its UFBoot2 support is reported as 0 (e.g., Anc-HeR in panel (C)). Full HeR+SzR+outgroup phylogeny, inferred with IQ-TREE from the MAFFT L-INS-i alignment, is shown in Fig. S1. Subset trees for SzR+outgroup and HeR+outgroup datasets are shown in Fig. S2.

### Indel-aware refinement restores compact, high-confidence ancestral rhodopsins

Indel treatment emerged as a major source of uncertainty in full-length ASR. When ancestral states were inferred in IQ-TREE without explicit indel modeling, the resulting ancestral sequences expanded far beyond extant rhodopsins, with the inflation concentrated in extra-membrane regions (Fig. 2A and Table 3). Under the PSI/TM-Coffee alignment, Anc-SzR, Anc-HeR, and Anc-SH reached 401, 434, and 408 residues, respectively, compared with 202 residues for SzR4 and 256 residues for HeR 48C12. This over-extension depressed mean site-wise posterior probabilities (PP; ∼38–66%) and, upon AlphaFold evaluation, produced models that were globally 7TM-like but dominated by elongated, low-confidence tails and EM segments (mean pLDDT typically ∼60–70; computed from per-residue C*α* pLDDT; see Materials and Methods). The same qualitative behavior was observed with the MAFFT L-INS-i alignment, indicating that in this dataset gap-unaware ASR on untrimmed alignments tends to inflate EM regions and produce low-confidence termini (Table 3).

**Figure 2:**
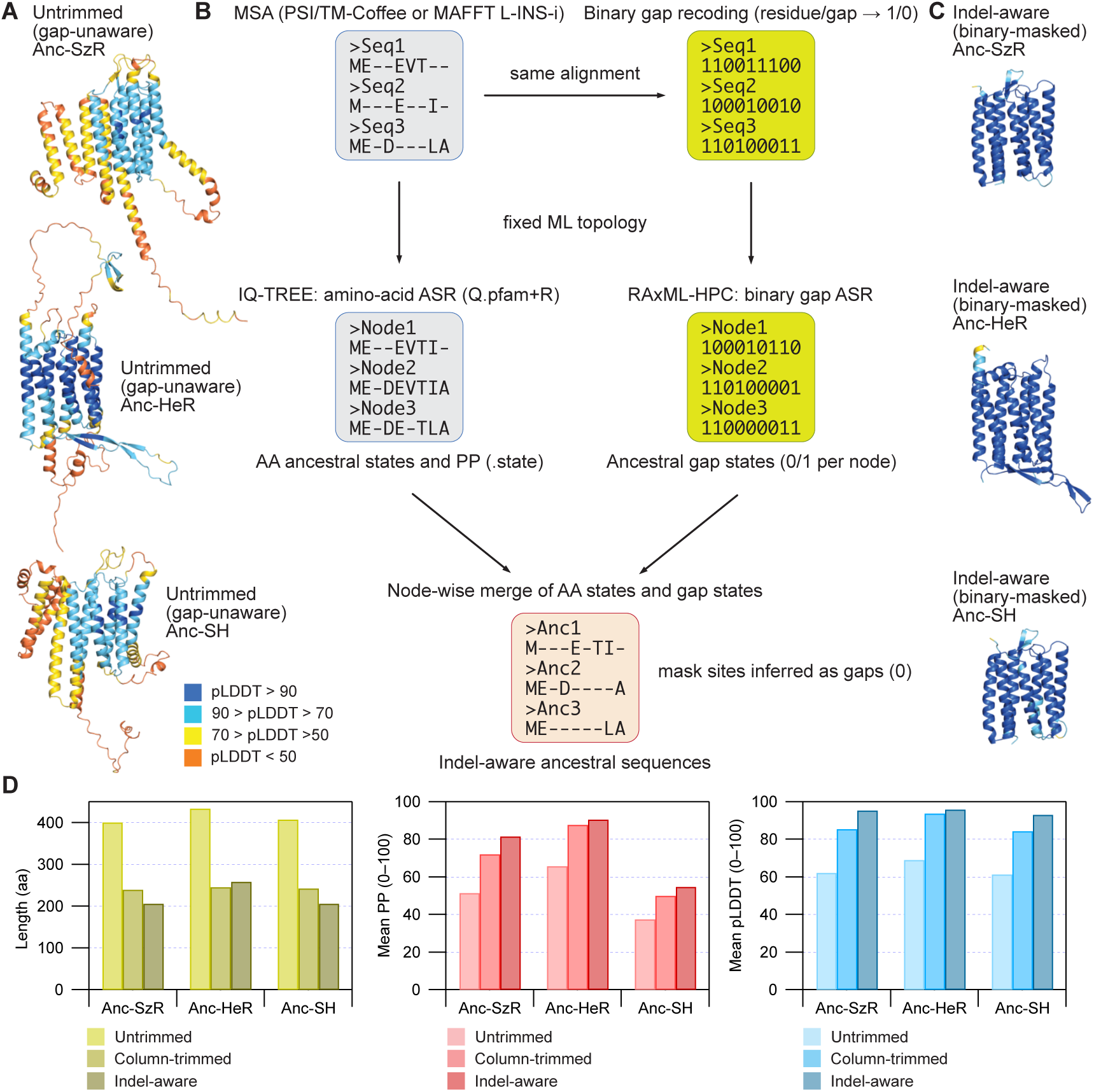
Indel-aware refinement restores realistic sequence lengths and improves statistical and structural confidence. (A) AlphaFold monomer models for untrimmed, gap-unaware ancestral sequences inferred directly from the full-length PSI/TM-Coffee alignment under Q.pfam+R7, without explicit indel treatment. Structures are colored by per-residue pLDDT using the standard AlphaFold color scale. All three ancestors form overall 7TM-like folds but exhibit overextended extra-membrane segments and low-confidence termini. (B) Schematic overview of the indel-aware ASR pipeline. The same full-length amino-acid alignment is used for amino-acid ASR in IQ-TREE and for binary gap coding followed by ancestral gap-state inference in RAxML on the fixed IQ-TREE topology. Node-specific gap reconstructions are then merged with the IQ-TREE .state files to mask sites inferred as gaps at each internal node, yielding indel-corrected ancestral sequences. (C) AlphaFold monomer models for the indel-corrected ancestors obtained after node-specific gap masking, colored as in panel (A). Indel-aware correction contracts overextended extra-membrane regions, removes low-confidence tails, and yields compact 7TM folds with uniformly high pLDDT. (D) Comparison of sequence length and confidence metrics for Anc-SzR, Anc-HeR, and Anc-SH under three reconstruction strategies: untrimmed gap-unaware ASR, a column-trimmed baseline using trimAl gappyout, and indel-aware refinement. Bar plots show sequence length, mean posterior probability (PP), and mean pLDDT. Mean pLDDT values were calculated from AlphaFold models, and mean PP and mean pLDDT are shown on a 0–100 scale. Relative to the untrimmed and column-trimmed baselines, indel-aware refinement yields more compact ancestors together with improved statistical and structural confidence (see Table 3). AlphaFold models for the column-trimmed baseline are shown in Fig. S3.

**Table 3:**
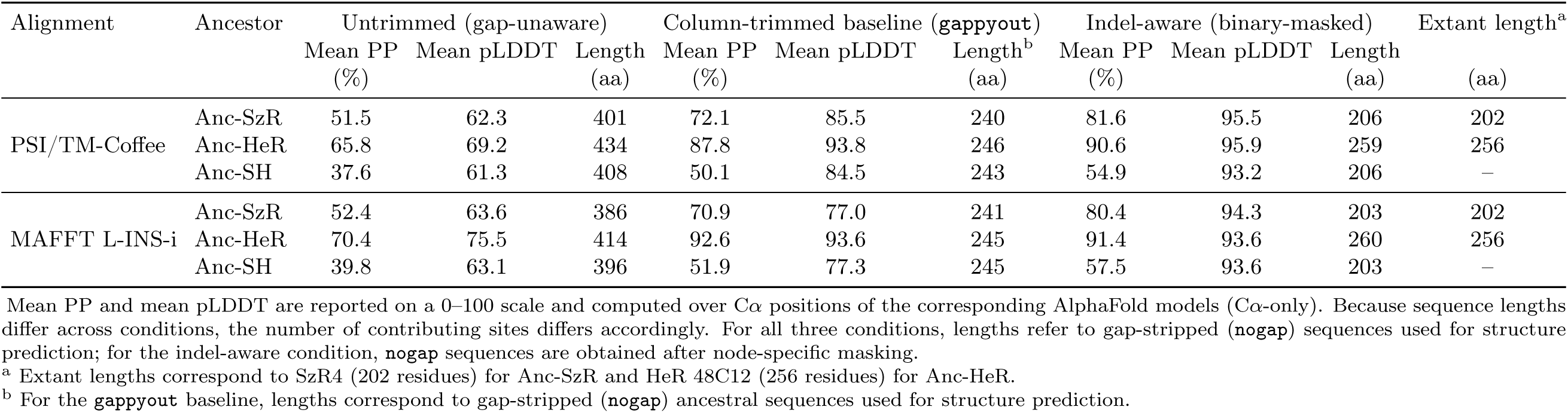
Global impact of indel-aware reconstruction on ancestral node statistics, with a column-trimming baseline.

To address this issue, a two-stage indel-aware refinement pipeline based on binary gap coding was implemented (*27*). Briefly, amino-acid ancestral states were inferred in IQ-TREE under profile-based models, while ancestral gap states were inferred from the same alignment recoded as a binary gap/non-gap matrix using a two-state model in RAxML on the fixed IQ-TREE topology. The inferred ancestral gap patterns were then mapped onto the IQ-TREE .state output, and sites inferred as gaps for each node were masked to generate node-specific, indel-corrected sequences (workflow summarized in Fig. 2B). This node-specific masking contracts each ancestor according to its inferred indel history, rather than applying a single global gap filter to all nodes.

Indel-aware correction had a pronounced impact on both statistical and structural metrics (Fig. 2C, D and Table 3). For the PSI/TM-Coffee alignment, mean PP increased from 51–66% to 82–91% for Anc-SzR and Anc-HeR, and mean pLDDT rose from ∼62–69 to *>*93. In parallel, sequence lengths contracted to values closely matching extant structures: 206 residues for Anc-SzR and 259 residues for Anc-HeR, compared with 202 residues for SzR4 and 256 residues for HeR 48C12. Comparable improvements were obtained with the MAFFT L-INS-i alignment, with Anc-SzR and Anc-HeR contracting to 203 and 260 residues, respectively, indicating that the qualitative effect of indel-aware refinement is reproducible across alignment strategies.

As a conventional column-trimmed baseline, we also performed gap-unaware ASR on column-trimmed alignments generated with trimAl gappyout. Column trimming reduced the most extreme over-extension and improved mean PP and pLDDT relative to untrimmed, gap-unaware ASR; however, it did not consistently recover compact, extant-like ancestors. For example, under PSI/TM-Coffee, gappyout yielded ancestors of 240–246 residues with mean pLDDT 85–94, whereas indel-aware masking produced 206 (Anc-SzR) and 259 (Anc-HeR) residues with mean pLDDT ∼96 (Table 3). Notably, the gappyout baseline forces a single global column filter across all nodes, whereas indel-aware masking yields node-specific lengths that track the inferred indel history (Table 3). These results indicate that global column trimming can partially mitigate gap-driven artifacts, whereas explicit indel-aware refinement more reliably restores compact ancestors from untrimmed alignments. Thus, the improvement is not only due to removing ambiguous columns globally, but arises from node-specific indel histories inferred on the same topology. Representative AlphaFold models for the column-trimmed baseline are shown in Fig. S3.

AlphaFold models of the indel-corrected ancestors formed compact 7TM folds with uniformly high pLDDT and retained lineage-specific extra-membrane motifs, including the short *β*-strands between TM2 and TM3 in SzR4 and the combination of a short *α*-helix between TM2 and TM3 with long *β*-strands between TM1 and TM2 in HeR 48C12 (*23*, *24*). In contrast, models predicted from uncorrected ancestors exhibited extended, low-confidence extra-membrane tails. Thus, node-specific masking based on the inferred ancestral indel state is a key step for obtaining full-length 7TM ancestors that are both statistically well supported and structurally coherent. All three ancestors retained the retinal-binding lysine in TM7; the FSE motif in TM3 was conserved in Anc-SzR but shifted to DSE in Anc-SH.

Beyond global statistics, extra-membrane (EM) secondary-structure patterns between TM1–TM2 and TM2–TM3 provide a concrete example of lineage-specific architectural change captured by the indel-aware reconstructions (Fig. 3). In extant structures, SzR4 displays short *β*-strands between TM2 and TM3 and lacks extended secondary structure between TM1 and TM2, whereas HeR 48C12 features a short *α*-helix between TM2 and TM3 together with long *β*-strands between TM1 and TM2 (Fig. 3A, B). The indel-corrected ancestors recapitulate these motifs in a clade-specific manner: Anc-SzR retains SzR4-like short *β*-strands between TM2 and TM3 and a loop-like TM1–TM2 region, whereas Anc-HeR retains the HeR-like combination of a TM2–TM3 short helix and TM1–TM2 long *β*-strands. By contrast, Anc-SH preserves only the SzR-like TM2–TM3 *β*-strands and lacks the HeR-specific helix/long-*β* combination, consistent with acquisition of these EM elements along the HeR lineage after divergence from Anc-SH (Fig. 3C). Consequently, indel-aware, node-specific masking can help retain informative EM architecture and enables gain/loss of secondary-structure motifs to be traced directly across ancestral nodes.

**Figure 3:**
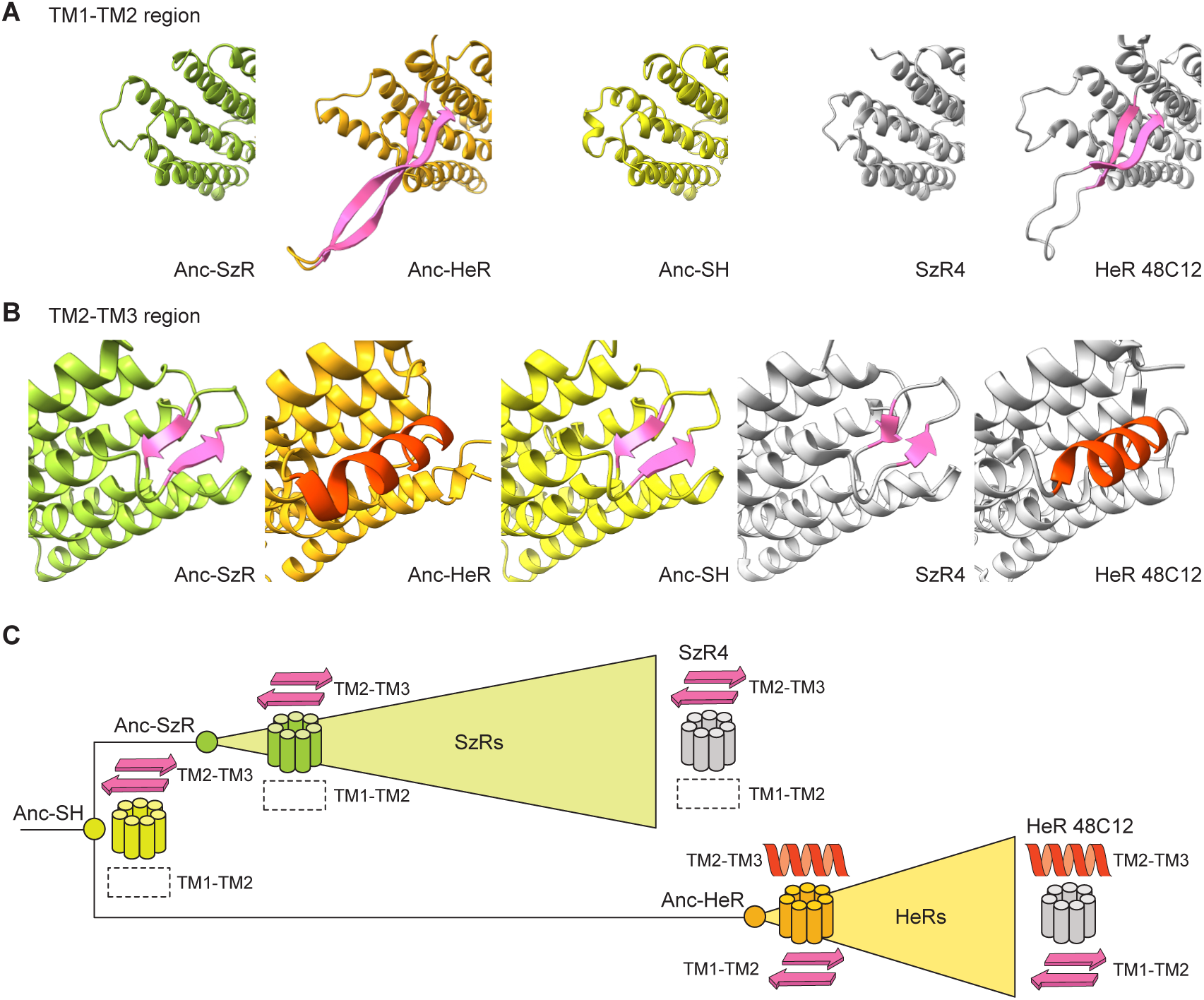
Evolutionary gain and loss of extra-membrane secondary-structure motifs between TM1–TM2 and TM2–TM3. (A) Close-up views of the extra-membrane segment between TM1 and TM2 for indel-corrected Anc-SzR (green), Anc-HeR (orange), Anc-SH (light green), and the extant structures SzR4 (PDB 7e4g) and HeR 48C12 (PDB 6su3; both light gray), aligned to their respective structural templates. *β*-strands in this region are highlighted in pink. Anc-HeR and HeR 48C12 share long *β*-strands between TM1 and TM2, whereas Anc-SzR, Anc-SH, and SzR4 lack defined secondary structure at this position and form a loop. (B) Corresponding close-up views of the extra-membrane segment between TM2 and TM3. *β*-strands are highlighted in pink and short *α*-helices in red. In Anc-SH, Anc-SzR, and SzR4, this region forms short *β*-strands, whereas in Anc-HeR and HeR 48C12 it forms a short *α*-helix, indicating a lineage-specific shift in secondary-structure motifs along the HeR branch. (C) Schematic summary of motif gain and loss mapped onto a simplified HeR–SzR phylogeny. Each ancestor or extant protein is represented by a seven-cylinder cartoon indicating the TM helices, with the TM2–TM3 and TM1–TM2 regions annotated. Pink arrows denote *β*-strands, red ribbons denote short *α*-helices, and dashed boxes mark loop-like regions lacking defined secondary structure. Together, these panels indicate that Anc-SzR and SzR4 largely retain the Anc-SH pattern (short *β*-strands between TM2 and TM3 and a loop between TM1 and TM2), whereas the HeR lineage loses the TM2–TM3 *β*-strands, gains a short *α*-helix in their place, and acquires long *β*-strands between TM1 and TM2.

We next quantified the similarity of indel-corrected ancestors across alignment strategies and relative to extant structural templates. PSI/TM-Coffee– and MAFFT L-INS-i–based reconstructions were highly consistent, with pairwise identities of 88.7% (Anc-SzR), 88.8% (Anc-HeR), and 82.8% (Anc-SH), suggesting convergence on similar solutions despite alignment differences. The indel-corrected ancestors also retained moderate similarity to extant rhodopsins: Anc-SzR (PSI/TM-Coffee and MAFFT L-INS-i) shared 54.0% identity with SzR4, and Anc-HeR shared 51.0% (PSI/TM-Coffee) and 47.1% (MAFFT L-INS-i) identity with HeR 48C12. These identity levels provide a quantitative context for the structural comparisons presented below.

### Region-level reliability across membrane-embedded and extra-membrane regions

To quantify region-level reliability, we partitioned each ancestral model into TM and EM regions using membrane boundaries from the Orientations of Proteins in Membranes (OPM) database (*41*). TM/EM boundaries follow the OPM membrane-embedding annotation. For SzR and HeR, we used the OPM entries for SzR4 and HeR 48C12, respectively. TM regions were defined as residues embedded within the OPM membrane slab (i.e., the membrane-embedded cores of the seven helices), whereas EM regions included all residues outside the slab, including helix termini, *β*-strands, and short coils. According to this convention, helical residues outside the membrane slab are classified as EM.

Using the structure-consistent alignments, OPM-derived TM/EM annotations were transferred from SzR4 and HeR 48C12 to the indel-corrected ancestors, and region-level reliability metrics were computed from PP and AlphaFold confidence (pLDDT) (Fig. 4A, Table 4, and Tables S5–S8). For each region, we summarized mean PP, mean pLDDT, and two combined indices (PP×pLDDT and PP−pLDDT). Anc-SH was not included in Table 4 because its hybrid SzR/HeR character precludes a single unambiguous template mapping; we therefore restrict Table 4 to the two terminal ancestors. However, its global trends are described qualitatively below.

**Figure 4:**
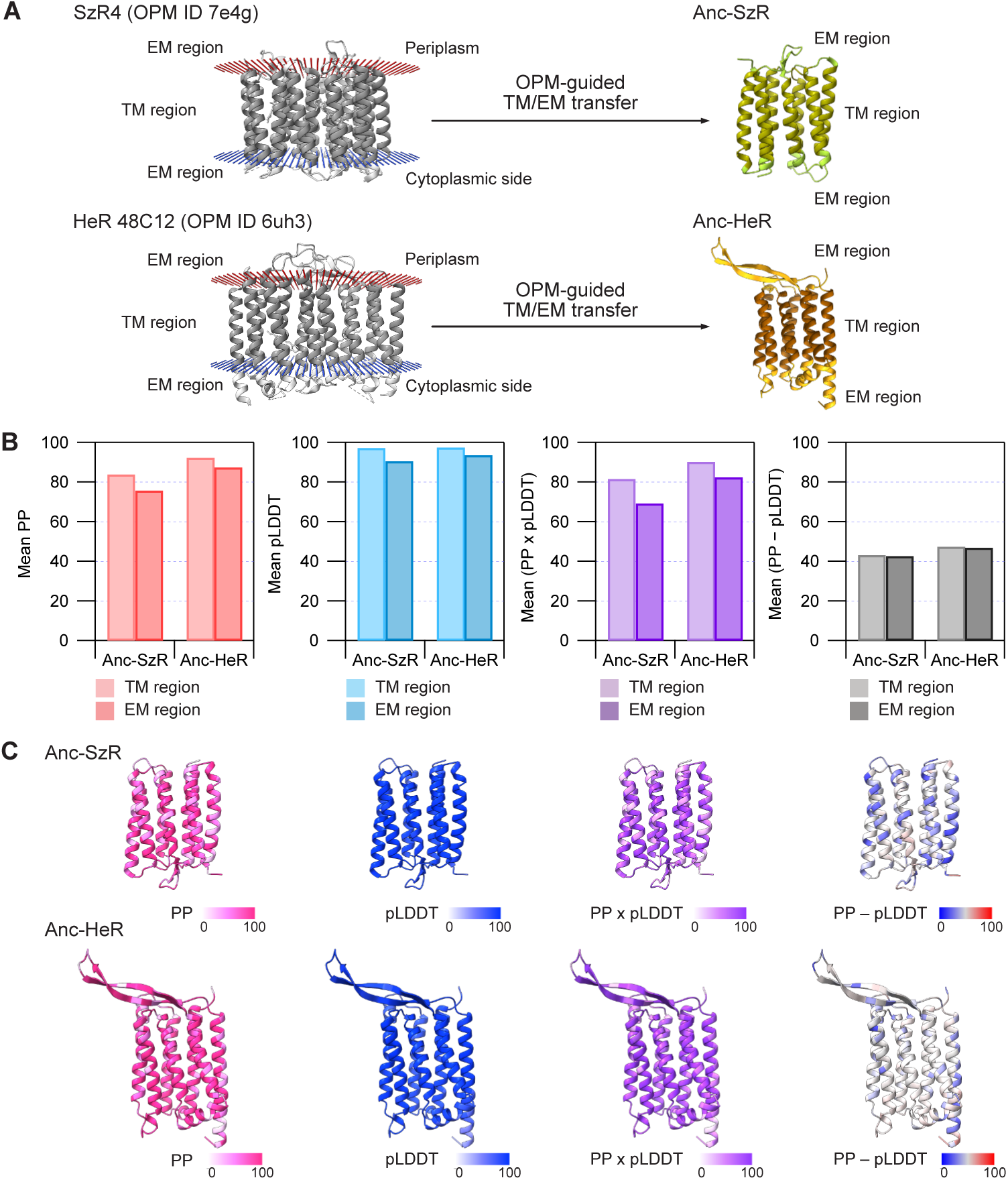
Region-level reliability of TM and EM segments in Anc-SzR and Anc-HeR. (A) OPM-guided transfer of transmembrane (TM) and extra-membrane (EM) annotations from extant structures to indel-corrected ancestors. OPM-derived TM/EM boundaries for SzR4 and HeR 48C12 are projected onto Anc-SzR and Anc-HeR via the PSI/TM-Coffee alignment under Q.pfam+R7. In the ancestral models, TM residues are colored dark green (Anc-SzR) or brown (Anc-HeR), and EM residues light green (Anc-SzR) or orange (Anc-HeR). (B) Region-level summary statistics for Anc-SzR and Anc-HeR. For each ancestor, bar plots report mean PP, mean pLDDT, mean (PP×pLDDT), and mean (PP−pLDDT) separately for TM and EM regions defined in panel (A). All metrics are shown on a 0–100 scale. TM segments display uniformly high confidence, whereas EM regions show only slightly reduced but still high values, indicating that many extra-membrane segments are reliably reconstructed. (C) Per-residue reliability mapped onto Anc-SzR and Anc-HeR structures. For each ancestor, four views are shown with C*α* atoms colored by PP (white to red), pLDDT (white to blue), PP×pLDDT (white to purple), or PP−pLDDT (blue through gray to red, indicating negative to positive), using the same 0–100 scale. These maps highlight that TM cores are consistently high-confidence, whereas local pockets of reduced PP with high pLDDT in EM segments mark positions where the fold is well defined but residue identity remains more ambiguous. Per-residue reliability mapped onto Anc-SH is shown in Fig. S4.

**Table 4:** Region-level reliability of ancestral 7TM rhodopsins. Mean posterior probabilities (PP), AlphaFold confidence (pLDDT), and derived indices for transmembrane (TM) and extra-membrane (EM) segments.

| Alignment | Ancestor | Region | Mean PP (%) | Mean pLDDT | Mean (PP×pLDDT) | Mean (PP–pLDDT) |
| --- | --- | --- | --- | --- | --- | --- |
| PSI/TM-Coffee | Anc-SzR | TM | 83.9 | 97.4 | 81.7 | 43.2 |
|  |  | EM | 75.9 | 90.7 | 69.3 | 42.6 |
|  | Anc-HeR | TM | 92.5 | 97.6 | 90.3 | 47.4 |
|  |  | EM | 87.5 | 93.7 | 82.5 | 46.9 |
| MAFFT L-INS-i | Anc-SzR | TM | 82.1 | 96.2 | 79.2 | 42.9 |
|  |  | EM | 76.2 | 89.3 | 68.0 | 43.5 |
|  | Anc-HeR | TM | 93.6 | 95.5 | 89.5 | 49.1 |
|  |  | EM | 88.0 | 91.3 | 81.0 | 48.3 |

TM and EM regions were defined by mapping OPM-derived membrane boundaries for SzR4 and HeR 48C12 onto Anc-SzR and Anc-HeR (see Methods). Values are means over C $\alpha$ positions retained after indel-aware masking.

Three consistent patterns emerged. First, TM regions were uniformly high-confidence. Across both alignments, TM segments in Anc-SzR and Anc-HeR exhibited high mean PP (approximately 82–94%) together with near-ceiling mean pLDDT values (*>*95) (Fig. 4B, C and Table 4). Accordingly, PP×pLDDT was highest in TM regions, indicating strong agreement between sequence-based support and structure-based confidence in the membrane-embedded helix cores.

Second, EM regions also showed unexpectedly high reliability. Although EM values were modestly lower than TM on average, mean PP and pLDDT remained high, with EM segments typically showing mean PP values of ∼76–88% and mean pLDDT values of ∼89–94 (Fig. 4B, C and Table 4). PP×pLDDT likewise indicated that many extra-membrane residues are well supported despite alignment uncertainty. Notably, EM segments corresponding to the characteristic *β*-strand motifs in SzR4 and HeR 48C12 retained high PP and pLDDT in Anc-SzR and Anc-HeR, indicating that these lineage-specific EM architectures are recovered as structured elements rather than being erased by indel-aware refinement.

Third, localized reductions in support were concentrated in TM1 and in peri-membrane EM segments flanking TM5. Across ancestors and alignments, PP was consistently lower in these regions, mirrored by reduced PP×pLDDT in the corresponding helix/loop segments (Tables S5 and S7). These sites likely represent evolutionarily labile positions where fold-level constraints are maintained but residue identities vary more freely.

The PP−pLDDT difference provided complementary local insight. While region-averaged PP−pLDDT values were similar between TM and EM segments (Fig. 4B, C and Table 4), site-wise profiles separated two common scenarios: positions with low PP but high pLDDT, where the fold is well defined despite ambiguity in residue identity, and positions with high PP but lower pLDDT, which tend to occur in intrinsically flexible EM segments. In this way, PP−pLDDT helps distinguish sequence-level uncertainty from structural uncertainty at the residue scale.

These region-level analyses showed that OPM-guided TM/EM partitioning provides a more informative view than a simple TM-versus-loop dichotomy, and that the indel-corrected full-length ancestors preserve substantial, lineage-specific extra-membrane architecture.

### Node-level reliability: divergence between evolutionary and structural support

We next assessed node-level reliability for three focal ancestors: Anc-SzR, Anc-HeR, and their MRCA Anc-SH. For each node, we summarized (i) branch support from the UFBoot2 consensus tree for the corresponding split, (ii) mean posterior probability (PP) across sites in the indel-corrected ancestor, and (iii) mean pLDDT from the AlphaFold model (Table 2). When a split was not recovered in the UFBoot2 consensus (i.e., collapsed into a polytomy), we set its support to 0 to reflect unresolved topology. UFBoot2 and SH-aLRT values on the ML tree are shown in Fig. 1, and Table 2 reports both ML-tree support (SH-aLRT/UFBoot2) and the UFBoot2 consensus frequencies used in downstream summaries.

Anc-SzR showed consistently strong support across all metrics. In the UFBoot2 consensus trees, the Anc-SzR split was recovered with high frequency (99% under PSI/TM-Coffee; 96% under L-INS-i), and the indel-corrected Anc-SzR sequences had high mean PP (80–82%) and mean pLDDT (94–96). TM segments in particular reached near-ceiling pLDDT values together with high PP, consistent with a well-resolved split and a structurally coherent 7TM ancestor. Collectively, these metrics identify Anc-SzR as a robust target for experimental resurrection.

Anc-HeR showed a contrasting pattern: strong sequence/structure support but alignment-sensitive topological support. Under PSI/TM-Coffee, the Anc-HeR split appeared in the UFBoot2 consensus with only moderate support (54%), despite very high mean PP (∼91%) and mean pLDDT (∼96) for the indel-corrected Anc-HeR sequence. Under L-INS-i, the same split collapsed in the UFBoot2 consensus (treated as 0% support), again despite similarly high PP and pLDDT. This discrepancy indicates that bootstrap replicates frequently sample alternative, near-equivalent arrangements within the HeR subtree. Thus, Anc-HeR is well supported conditional on a chosen local topology, but the local HeR branching order remains sensitive to alignment details.

Anc-SH showed the opposite trade-off. The deeper split separating HeRs and SzRs from the outgroup had maximal UFBoot2 support (100%) across all conditions, indicating a robust topology. In contrast, the indel-corrected Anc-SH sequences had substantially lower mean PP (∼55–57%) than the terminal ancestors, even though mean pLDDT remained high (∼93–94) and the overall 7TM fold was well formed. Thus, the deep split is stable, but residue identities at Anc-SH are more diffuse, with many sites supported by multiple plausible amino-acid states.

To assess UFBoot2 convergence, we repeated the bootstrap analyses with 1,000, 2,000, and 5,000 replicates and, for the most demanding setting, an independent run with an alternative random seed (Table S4). Because UFBoot2 relies on stochastic resampling and heuristic tree searches, consensus frequencies can vary modestly across runs, especially for locally unstable regions of the tree. Under PSI/TM-Coffee, the overall interpretation was stable: Anc-SzR and Anc-SH remained consistently high (97–99% and 100%, respectively), whereas Anc-HeR showed moderate but non-maximal support. Importantly, the baseline value reported in Table 2 (54%) is consistent with the moderate range observed across repeated analyses (54–64%; Table S4). Under MAFFT L-INS-i, Anc-SzR and Anc-SH again remained stable (95–98% and 100%), whereas Anc-HeR remained highly sensitive and ranged from being absent in the UFBoot2 consensus (collapsed into a local polytomy) to moderate support (47%) at 5,000 replicates (Table S4), consistent with the local instability described above.

Together, these contrasts demonstrate that no single metric—branch support, PP, or pLDDT—adequately captures ASR reliability. Stable topology can coexist with diffuse residue support, and high PP/pLDDT can occur despite alignment-sensitive branching. Therefore, we introduce a composite index that integrates all three dimensions into a node-level reliability score.

### Extant sequence reconstruction supports quantitative accuracy of the baseline pipeline

As a direct validation of quantitative reconstruction accuracy, we performed an extant sequence reconstruction (ESR) proxy test (*40*). In this proxy design, a chosen target tip (SzR4 or HeR 48C12) is removed from the inference as a sequence and replaced by a proxy subtree comprising two dummy leaves (suffixes_A and _B) attached by long branches (dummy branch length = 500), while the internal proxy node retains the original target label. The corresponding alignment entry is replaced by two fixed dummy sequences (all L and all V). We then re-ran IQ-TREE ASR under the baseline setting (PSI/TM-Coffee; Q.pfam+R7) on the fixed proxy topology and scored, at sites where the true target residue was not a gap, (i) the identity between the maximum-posterior reconstructed residue and the true residue and (ii) the posterior probability assigned to the true residue (Table 5). Because the multiple sequence alignment was treated as fixed (i.e., not recomputed after removing the target), this ESR should be interpreted as a practical proxy that evaluates reconstruction accuracy conditional on the chosen alignment/model configuration.

**Table 5:**
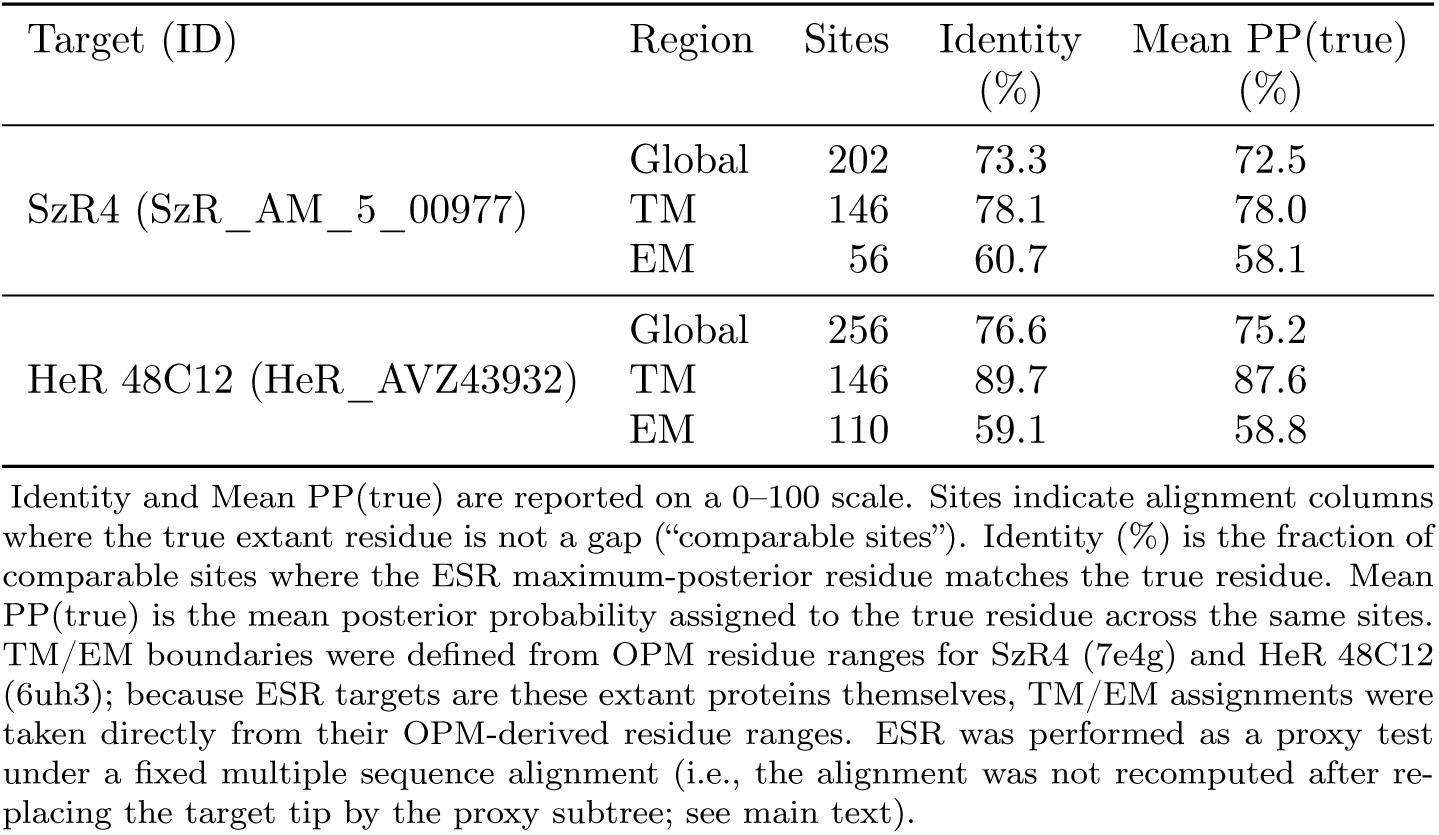
Extant sequence reconstruction (ESR) proxy validation under the base-line setting (PSI/TM-Coffee; Q.pfam+R7).

| Target (ID) | Region | Sites | Identity (%) | Mean PP(true) (%) |
| --- | --- | --- | --- | --- |
| SzR4 (SzR_AM_5_00977) | Global | 202 | 73.3 | 72.5 |
|  | TM | 146 | 78.1 | 78.0 |
|  | EM | 56 | 60.7 | 58.1 |
| HeR 48C12 (HeR_AVZ43932) | Global | 256 | 76.6 | 75.2 |
|  | TM | 146 | 89.7 | 87.6 |
|  | EM | 110 | 59.1 | 58.8 |

Identity and Mean PP(true) are reported on a 0–100 scale. Sites indicate alignment columns where the true extant residue is not a gap (“comparable sites”). Identity (%) is the fraction of comparable sites where the ESR maximum-posterior residue matches the true residue. Mean PP(true) is the mean posterior probability assigned to the true residue across the same sites. TM/EM boundaries were defined from OPM residue ranges for SzR4 (7e4g) and HeR 48C12 (6uh3); because ESR targets are these extant proteins themselves, TM/EM assignments were taken directly from their OPM-derived residue ranges. ESR was performed as a proxy test under a fixed multiple sequence alignment (i.e., the alignment was not recomputed after replacing the target tip by the proxy subtree; see main text).

For SzR4 (202 comparable sites), the ESR reconstruction achieved 73.3% global identity with a mean PP assigned to the true residue of 72.5%. Reconstruction accuracy was higher in TM regions than in EM regions: TM identity was 78.1% (meanPP(true) = 78.0%), whereas EM identity was 60.7% (meanPP(true) = 58.1%). For HeR 48C12 (256 comparable sites), ESR accuracy was similarly strong overall (76.6% global identity; meanPP(true) = 75.2%) and again showed a pronounced TM/EM contrast (TM: 89.7%, meanPP(true) = 87.6%; EM: 59.1%, meanPP(true) = 58.8%). Thus, under the baseline alignment/model configuration used throughout this study, the pipeline recovers held-out extant rhodopsins with high accuracy in TM cores and moderate accuracy in EM regions, consistent with the expected distribution of evolutionary constraint across structural compartments.

To assess whether these accuracies are specific to the two structural reference proteins, we additionally performed the same ESR proxy test for four additional SzR sequences and four additional HeR sequences and summarized global identity scores in Table S9. Across these additional targets, global ESR identities spanned a similar range (approximately 68–99% for SzR targets and 80–93% for HeR targets), indicating that the baseline ESR performance is not idiosyncratic to SzR4 and HeR 48C12.

### Composite reliability metric integrating PP, pLDDT, and BS

Building on this extant sequence validation, we return to the primary goal of this study: assessing the reliability of deep ancestral nodes whose true sequences are unknown. While ESR provides a quantitative baseline for reconstruction accuracy on extant targets, reliability at internal nodes additionally depends on local topological stability and on the consistency between sequence-level posterior support and structure-level confidence. We therefore introduce an explicit composite metric that integrates posterior probabilities (PP), AlphaFold confidence (pLDDT), and ultrafast bootstrap support (UFBoot2) to summarize node-level reliability in a single score.

To integrate evolutionary, statistical, and structural evidence at internal nodes, we defined a composite node-level reliability metric. We first quantified sequence–structure consistency as the mean per-site product 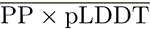, which is high only when both the posterior probability of the inferred residue and the predicted structural confidence at that site are high. We then modulated this value by UFBoot2 support for the corresponding split, measured as its frequency in the bootstrap consensus tree in the bootstrap consensus tree, to obtain

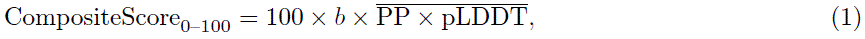

where *b* = UFBoot2*/*100 is the UFBoot2 support expressed as a fraction (0–1), and PP and pLDDT are each scaled to 0–1 before taking their product and averaging across sites. When a split was absent from the UFBoot2 consensus (collapsed into a polytomy), we set *b* = 0. Composite scores were reported on a 0–100 scale for interpretability (Table 2).

Applying this metric highlights a clear contrast: Anc-SzR is consistently robust, Anc-SH is intermediate, and Anc-HeR is strongly dependent on local topology. For PSI/TM-Coffee, mean PP×pLDDT values were 78, 87, and 52 for Anc-SzR, Anc-HeR, and Anc-SH, respectively, while the corresponding UFBoot2 supports were 99%, 54%, and 100% (Fig. 5A and Table 2), yielding composite scores of 77, 47, and 52. Under MAFFT L-INS-i, mean PP×pLDDT values were similar (76, 86, and 54), but UFBoot2 supports were 96%, 0%, and 100%, giving composite scores of 73 for Anc-SzR, 0 for Anc-HeR (the focal split is absent from the UFBoot2 consensus tree and is treated as unresolved), and 54 for Anc-SH (Fig. 5B and Table 2). Thus, Anc-SzR remains high across alignments, Anc-SH remains moderate, whereas Anc-HeR collapses when the HeR split is not reproducibly recovered.

**Figure 5:**
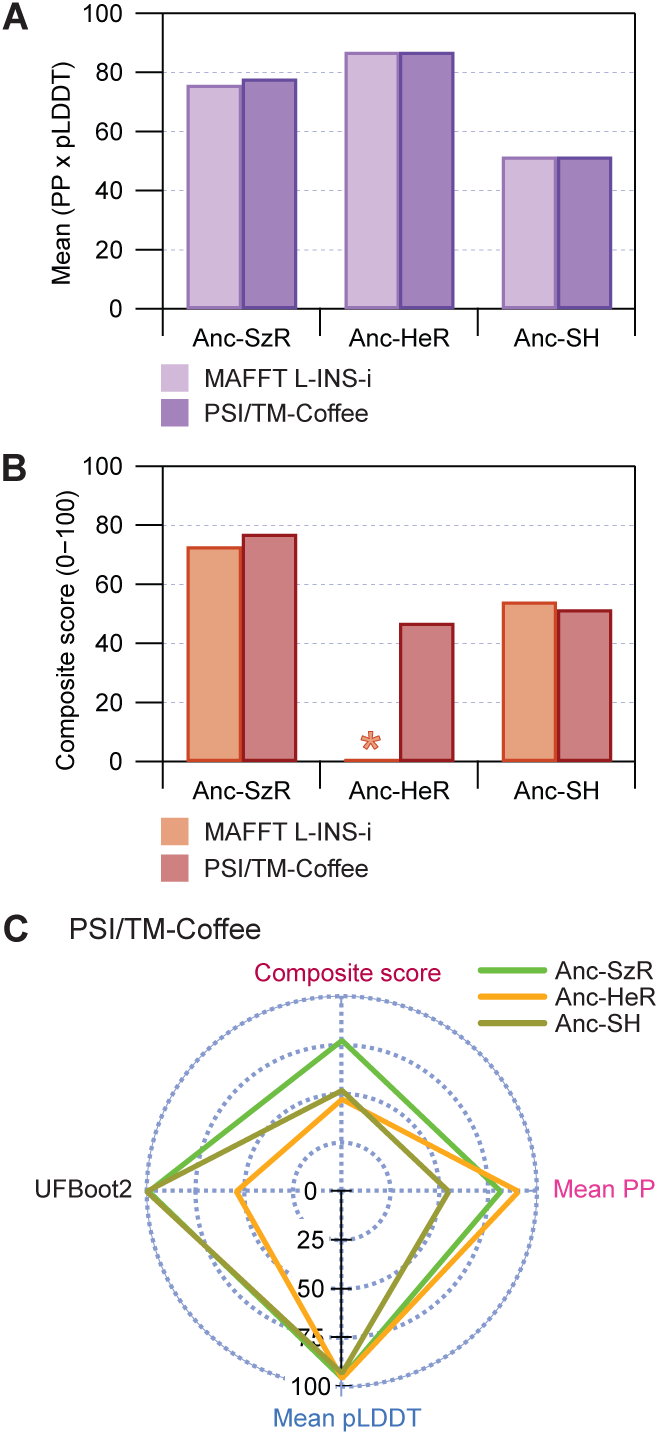
Node-level sequence–structure consistency and composite reliability across alignments. (A) Mean PP×pLDDT for Anc-SzR, Anc-HeR and Anc-SH under the PSI/TM-Coffee and MAFFT L-INS-i alignments. Values are averaged over C*α* sites and shown on a 0–100 scale. For all three nodes, mean PP×pLDDT is high and broadly similar between alignments, indicating strong overall sequence–structure consistency in the reconstructed ancestors. (B) Composite node-level reliability scores (0–100) combining UFBoot2 branch support and mean PP×pLDDT for the same nodes and alignments. The composite score is defined as 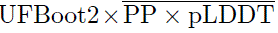 (see Methods). For the MAFFT L-INS-i alignment, the Anc-HeR split is not recovered in the UFBoot2 consensus (collapsed branch; UFBoot2 = 0; its composite score is therefore 0 and is marked with an asterisk. Across both alignments, Anc-SzR attains consistently high composite scores, Anc-SH reaches intermediate values, and Anc-HeR is conditionally reliable, with a high composite score only when the HeR split is recovered in the consensus. (C) Radar plot summarizing node-level metrics for the PSI/TM-Coffee alignment. Anc-SzR (green), Anc-HeR (orange) and Anc-SH (dark yellow) are shown with axes corresponding to mean PP, mean pLDDT, mean PP×pLDDT and the composite score (all on a 0–100 scale). The plot highlights that Anc-SzR is uniformly strong across all metrics, Anc-HeR combines very high sequence–structure consistency with alignment-sensitive branch support, and Anc-SH exhibits robust topology but lower residue-level certainty.

This decomposition clarifies why the same dataset can yield “confident” ancestors for different reasons. Anc-HeR combines very high PP×pLDDT with topology-sensitive UFBoot2; accordingly, its composite score is high under PSI/TM-Coffee but strongly suppressed under L-INS-i, matching the local instability described above. By contrast, Anc-SzR pairs high PP×pLDDT with consistently strong UFBoot2 support under both alignments, making it the most conservative target for experimental resurrection. Anc-SH shows the converse: consistently maximal UFBoot2 support but lower PP×pLDDT, underscoring that deep nodes can be topologically stable yet residue-wise diffuse (Fig. 5C).

We emphasize that this composite score is a decision-support summary rather than a substitute for inspecting BS, PP, and pLDDT individually. Nodes that are collapsed or extensively rearranged in the UFBoot2 consensus—or whose support is clearly alignment-dependent—should be flagged explicitly, and their composite scores interpreted as conditional. In this sense, Anc-HeR represents a conditionally reliable ancestor: its site-wise and structural confidence are strong, but its local evolutionary context remains sensitive to alignment choice.

### Oligomeric-state prediction supports dimeric HeR and trimeric SzR ancestors

To test whether the resurrected ancestors encode biologically plausible quaternary structure, we predicted homooligomeric assemblies and asked whether the preferred stoichiometries match those observed for extant rhodopsins. Using indel-corrected sequences inferred from the PSI/TM-Coffee alignment, we modeled Anc-SzR, Anc-HeR, and Anc-SH as dimers, trimers, and pentamers with an AlphaFold multimer workflow (Fig. 6). We focused on stoichiometries of 2, 3, and 5 because microbial rhodopsins are most commonly reported as dimers, trimers, or pentamers, whereas tetrameric assemblies are not well established for this family (*21*, *46*). For each target and stoichiometry, we recorded the predicted TM-score (pTM) and interface predicted TM-score (ipTM) as summaries of overall fold confidence and interface quality, respectively (*47*). Results for both alignment strategies (PSI/TM-Coffee and MAFFT L-INS-i) are summarized in Table S10.

**Figure 6:**
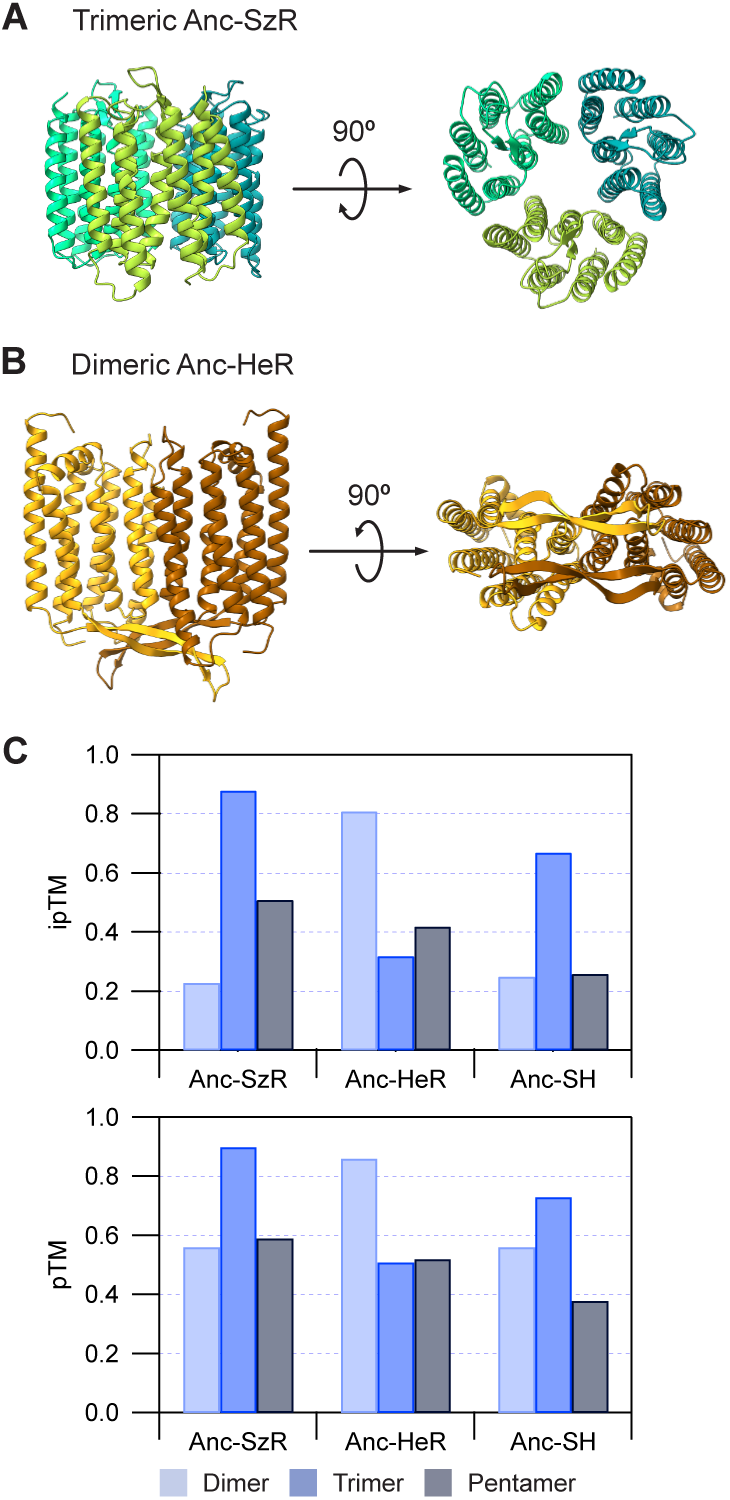
Oligomeric-state predictions support trimeric Anc-SzR and dimeric Anc-HeR. (A) AlphaFold-Multimer prediction of trimeric Anc-SzR viewed parallel to the membrane (left) and from the cytoplasmic side (right). Each protomer is shown as a cartoon and colored by chain. The predicted assembly forms a compact trimeric arrangement of seven-helix bundles. (B) AlphaFold-Multimer prediction of dimeric Anc-HeR viewed parallel to the membrane (left) and from the extracellular side (right), with protomers colored by chain. The predicted interface includes extra-membrane elements between TM1 and TM2, consistent with the location of proposed dimerization motifs in extant HeRs. (C) Predicted interface TM-scores (ipTM) and overall complex TM-scores (pTM) for dimeric, trimeric and pentameric assemblies of Anc-SzR, Anc-HeR and Anc-SH, based on indel-corrected sequences inferred from the PSI/TM-Coffee alignment. Bars show the top-ranked AlphaFold-Multimer model for each stoichiometry. Anc-SzR attains the highest ipTM/pTM as a trimer, Anc-HeR as a dimer, and Anc-SH shows intermediate ipTM values with a modest preference for trimeric assemblies. Tetrameric assemblies were not evaluated, as microbial rhodopsins are predominantly reported as dimers, trimers or pentamers rather than tetramers. The same qualitative ranking of preferred stoichiometries was observed for L-INS-i–based ancestors (Table S10).

Anc-SzR was strongly predicted to form a trimer (Fig. 6A, C). Under PSI/TM-Coffee, the trimer achieved high confidence (ipTM = 0.88; pTM = 0.90), whereas dimeric and pentameric assemblies scored substantially lower; MAFFT L-INS-i produced the same ranking (Table S10). This preference mirrors the trimeric assemblies reported for extant SzRs and suggests that both the 7TM core and the reconstructed EM elements support a plausible trimeric ancestral interface.

In contrast, Anc-HeR was strongly predicted to form a dimer (Fig. 6B, C). Under PSI/TM-Coffee, the dimer achieved high confidence (ipTM = 0.81; pTM = 0.86), while higher-order assemblies were markedly weaker; MAFFT L-INS-i yielded the same ranking (Table S10). Consistent with this, the dimeric models suggest an interface involving the EM *β*-strand region between TM1 and TM2, in line with the proposed role of this motif in extant HeRs.

The deeper Anc-SH showed intermediate behavior: trimeric models (ipTM = 0.67; pTM = 0.73) were preferred over dimers and pentamers, but with lower interface confidence than the terminal ancestors (Fig. 6C and Table S10), consistent with Anc-SH’s deeper phylogenetic position, lower mean PP, and mixed HeR/SzR character. Overall, these multimeric predictions support the conclusion that the indel-corrected ancestors encode oligomerization interfaces compatible with the trimeric and dimeric states observed for SzRs and HeRs, respectively.

### Expression and spectral characterization of ancestral rhodopsins

To evaluate the ability of reconstructed ancestral rhodopsins to be expressed and bind to retinal, we focused on Anc-SzR and Anc-HeR and expressed them heterologously in *E. coli* under multiple induction conditions. Importantly, the constructs were designed to use the ASR-derived sequences essentially “as reconstructed”. For each protein, the inferred ancestral sequence (PSI/TM-Coffee reconstruction) was cloned as a single open reading frame into a standard pET vector, with only a vector-encoded His_6_ affinity tag. Anc-SzR was inserted into pET-21a(+) with a C-terminal His_6_ tag, whereas Anc-HeR was inserted into pET-15b with an N-terminal His_6_ tag and a thrombin recognition site upstream of the ancestral open reading frame. No additional fusion partners, solubility tags, truncations, or stabilizing mutations were introduced. Thus, any rhodopsin-like pigmentation and spectral features observed in *E. coli* directly reflect the intrinsic properties of the full-length ancestral sequences produced by our pipeline.

For both Anc-SzR and Anc-HeR, we compared three induction regimes after addition of IPTG and all-*trans*-retinal: 37 °C for 4 h, 37 °C for 20 h, and 25 °C for 20 h. Under all conditions tested, harvested cell pellets displayed a visible reddish/purple coloration, consistent with expression of retinal-binding rhodopsins. Pigmentation was strongest after 20 h induction (both at 37 °C and 25 °C) and weaker after 4 h at 37 °C. We therefore adopted 25 °C for 20 h as the standard condition for subsequent preparations of both ancestors. Under this condition, both constructs reproducibly yielded strongly pigmented pellets (Fig. 7A), indicating efficient folding and chromophore incorporation in vivo without engineering beyond a simple His_6_ tag.

**Figure 7:**
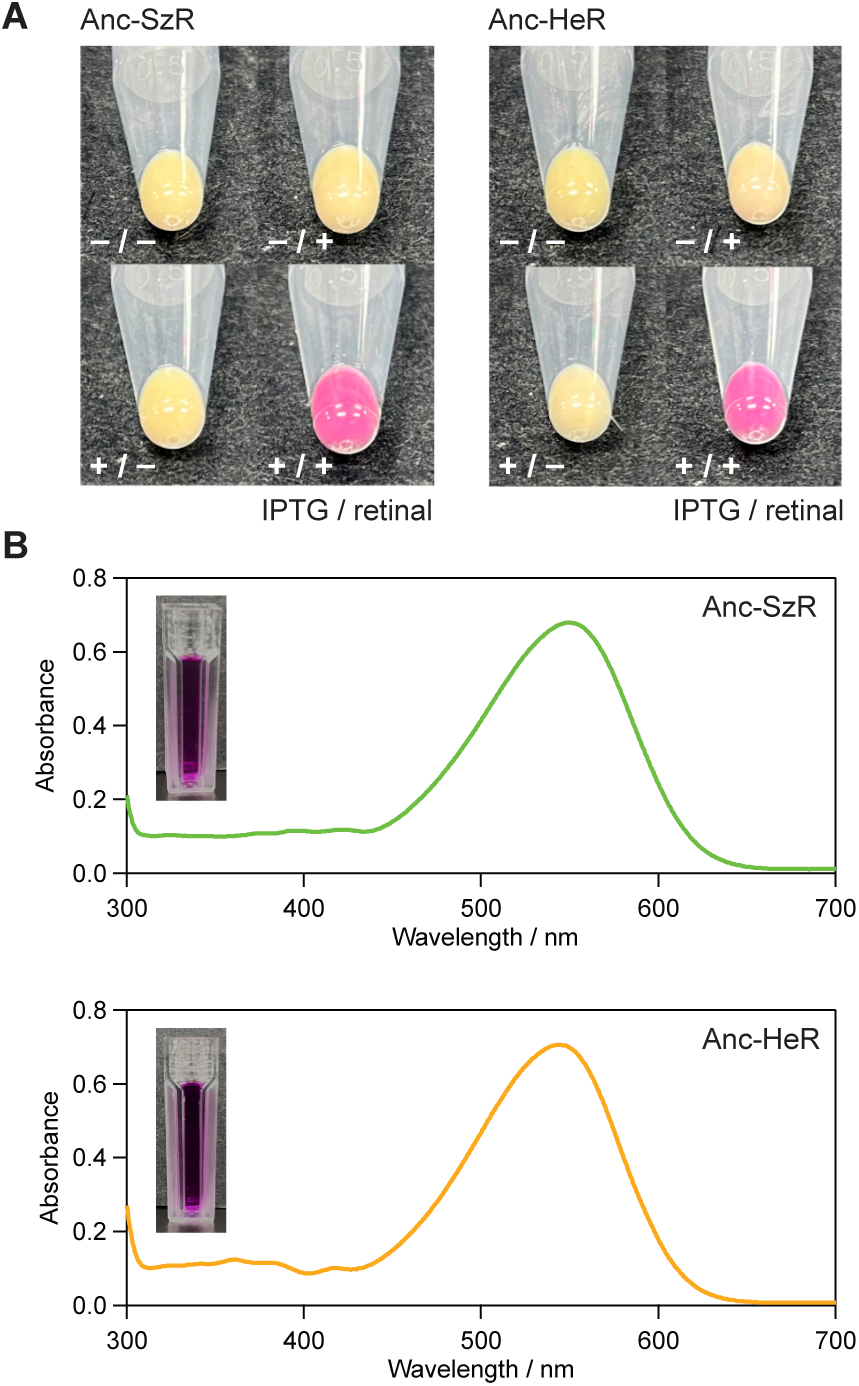
Expression and spectral properties of Anc-SzR and Anc-HeR. (A) Whole-cell pigmentation of *E. coli* expressing Anc-SzR or Anc-HeR under four combinations of IPTG induction and all-*trans*-retinal supplementation (−*/*−, −*/*+, +*/*−, +*/*+). Pronounced coloration is observed in induced cells supplied with retinal (+*/*+), whereas non-induced and/or retinal-free controls show minimal coloration, consistent with retinal-dependent holoprotein formation. (B) UV–visible absorption spectra of purified, detergent-solubilized Anc-SzR and Anc-HeR recorded at room temperature after affinity purification. Insets show photographs of the purified samples in cuvettes, illustrating the characteristic visible colors of the ancestral rhodopsins.

**Figure 8:**
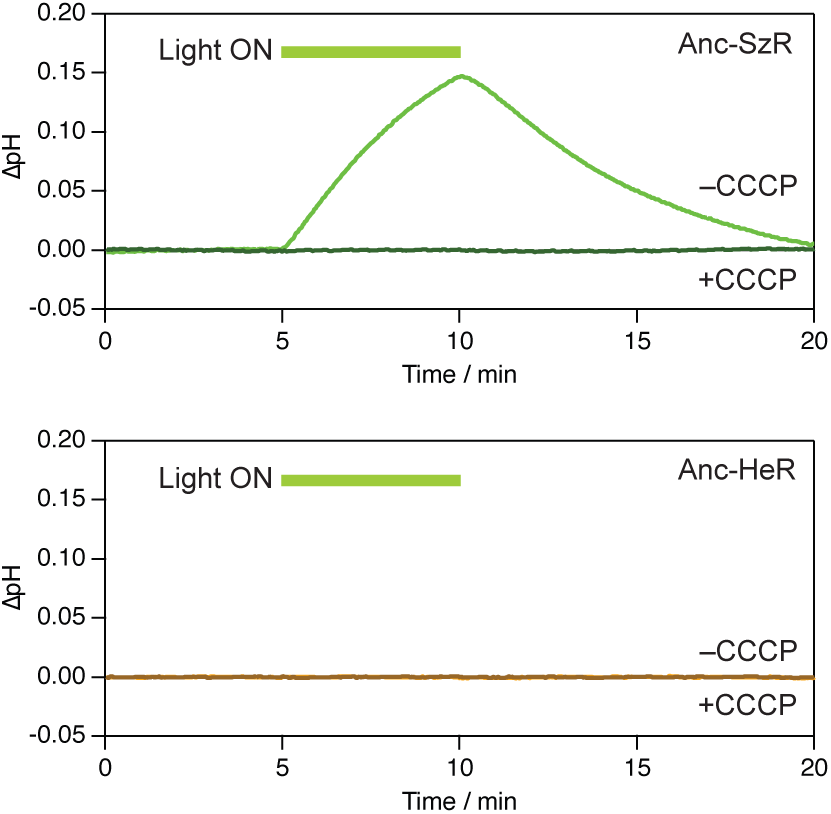
Whole-cell pH assay for Anc-SzR and Anc-HeR. External pH was monitored for unbuffered *E. coli* suspensions (300 mM NaCl) expressing Anc-SzR or Anc-HeR during during dark/light measurements; a representative single cycle is shown here, and raw two-cycle traces are provided in Fig. S6. Traces are shown as ΔpH after baseline subtraction (see Supplementary Methods); raw pH traces are provided in Fig. S6. Thick bars above the traces indicate illumination periods (540 nm). Traces are shown for Anc-SzR without CCCP (green) and with CCCP (30 µM; dark green), and for Anc-HeR without CCCP (orange) and with CCCP (dark orange). Anc-SzR shows a clear light-induced alkalinization (pH increase) that is largely abolished by CCCP, consistent with light-driven proton-transport activity. Anc-HeR shows no clear light-dependent pH change with or without CCCP within the detection limit of this whole-cell assay; this result is consistent with the prevailing view that extant heliorhodopsins are typically not canonical light-driven ion pumps.

Detergent-solubilized Anc-SzR and Anc-HeR were further assessed by UV–visible absorption spectroscopy. Purified Anc-SzR exhibited a dominant absorption maximum at 549 nm, whereas purified Anc-HeR showed a *λ*_max_ at 543 nm (Fig. 7B). These maxima are within the ranges reported for extant schizorhodopsins and heliorhodopsins, respectively (*20*, *22*), supporting that each reconstructed ancestor forms a retinal-binding holoprotein with clade-typical spectral properties. The overall spectral line shapes were also characteristic of a protein-bound retinal chromophore in an ordered environment, consistent with the compact, high-confidence 7TM folds predicted by AlphaFold. Photographs of the purified samples (Fig. 7B, inset) further illustrate the characteristic visible coloration of the ancestral rhodopsins.

### Light-driven proton transport activity in whole cells

To test whether the resurrected ancestors exhibit light-driven ion-transport activity beyond retinal binding, we performed a whole-cell pH assay commonly used for microbial rhodopsins (*42*). *E. coli* suspensions expressing Anc-SzR or Anc-HeR were resuspended in unbuffered 300 mM NaCl and subjected to repeated dark/light cycles (illumination at 540 nm), while monitoring external pH (Fig. 8). In the absence of CCCP, illumination of Anc-SzR-expressing cells induced a clear alkalinization of the external medium (pH increase) that was reproducible across on/off cycles within the same preparation. This light-driven pH response was largely abolished upon addition of CCCP (30 µM), consistent with light-driven proton-transport activity rather than secondary pH changes caused by transport of other ions or nonspecific photoeffects. In contrast, Anc-HeR-expressing cells showed no detectable light-driven pH change under the same conditions, with or without CCCP. This negative result is consistent with the prevailing view that heliorhodopsins are not canonical light-driven ion pumps, and it supports that Anc-HeR behaves in a manner broadly aligned with extant HeRs under this assay. Together, these results provide initial functional evidence that Anc-SzR supports light-driven proton transport under the conditions tested, while Anc-HeR shows no detectable ion-pumping activity in this whole-cell assay—consistent with the prevailing view for extant heliorhodopsins (*20*, *21*). Proton pumping has been reported for a giant-virus heliorhodopsin, but viral heliorhodopsins were not intentionally included in our dataset and thus do not affect the present Anc-HeR reconstruction (*31*).

## Discussion

This study provides a full-length case study of ASR in 7TM microbial rhodopsins, focusing on SzRs and HeRs. By combining structure-consistent alignments, profile-based evolutionary models, explicit indel-aware refinement, and structure-based evaluation, we reconstructed compact full-length ancestors (Anc-SzR, Anc-HeR, and Anc-SH) and demonstrated experimental tractability by expressing Anc-SzR and Anc-HeR in *E. coli* as stable, colored holoproteins with rhodopsin-like absorption maxima, and we further showed that Anc-SzR supports light-driven proton transport in a whole-cell pH assay. Together, these results show that the resurrection of full-length 7TM ancestors is feasible in this SzR–HeR system without stabilizing engineering and that EM architecture can be reconstructed and evaluated rather than routinely trimmed away.

A central practical finding from the HeR–SzR system is that alignment quality can dominate local phylogenetic stability, especially within the HeR clade. Across datasets and conditions, Q.pfam+R models were consistently preferred over conventional single-matrix models (Table 1), supporting the use of membrane-aware profile models for these sequences. However, the strongest qualitative differences we observed were not between Q.pfam+R and alternative amino-acid models, but between alignment strategies: PSI/TM-Coffee preserved TM registers and produced a HeR subtree that was more consistently recovered in UFBoot2 consensus trees. In contrast, MAFFT L-INS-i yielded a locally unstable resolution around the HeR ancestor (Fig. 1). In other words, for this case study, model choice improved global fit, but alignment quality governed whether key internal HeR splits were repeatedly recovered under resampling.

A second key finding is that explicit indel-aware refinement is essential for realistic full-length ancestors in this system. Gap-unaware ASR on full-length alignments produced markedly overextended ancestors with low-confidence tails in EM regions (Fig. 2A), a failure mode that is easy to miss if one focuses only on the TM core. Indel-aware refinement, implemented here by combining amino-acid ASR from IQ-TREE with node-specific binary gap inference in RAxML on a fixed topology, restored lengths that closely match extant structures and simultaneously increased both PPs and pLDDT in the retained sites (Fig. 2C, D). This “before/after” contrast is especially informative for 7TM proteins because termini and EM motifs can contribute to folding, oligomerization, and function; in the SzR–HeR case, indel-aware correction converts diffuse, elongated ancestors into compact sequences whose 7TM folds and EM motifs are predicted with uniformly high confidence. Importantly, this improvement was not replicated by a conventional column-trimming baseline (gappyout), which reduced over-extension but did not consistently recover compact, extant-like ancestors (Table 3 and Fig. S3).

Within this full-length framework, the EM regions of SzRs and HeRs proved more interpretable than commonly assumed. Using OPM-defined membrane boundaries to partition membrane-embedded TM cores from all residues outside the bilayer, we found that EM segments retained high mean PP and high mean pLDDT, only modestly reduced relative to TM regions (Fig. 4 and Table 4). ESR proxy validation on SzR4 and HeR 48C12, which provided a quantitative baseline for reconstruction accuracy and confirmed the expected TM>EM contrast in recoverability (Table 5). Importantly, EM secondary-structure motifs were recovered in a lineage-specific manner rather than being erased by indel-aware correction. In HeR, the long *β*-strands between TM1 and TM2 and the short helix between TM2 and TM3 seen in HeR 48C12 were recapitulated in Anc-HeR, whereas SzR-like ancestors retained short *β*-strands between TM2 and TM3 and a loop-like TM1–TM2 region (Fig. 3A, B). In this case study, these motifs behave as reconstructable evolutionary characters: Anc-SH retains the SzR-like pattern, while the HeR lineage shows a gain/loss and relocation of EM elements along the branch to Anc-HeR (Fig. 3C). This provides a concrete example of how full-length ASR enables evolutionary statements about EM architecture rather than restricting inference to the TM core alone.

By comparing node-level metrics, we also highlight that topological support, residue-level certainty, and structural confidence can diverge in ways that matter for experimental prioritization. Anc-SzR is robust across alignments, with high UFBoot2 support, high mean PP, and high mean pLDDT, making it a conservative resurrection target (Fig. 1 and Table 2). Anc-HeR, by contrast, shows very high PP and pLDDT once a particular local HeR topology is chosen, but its surrounding subtree is alignment-sensitive, leading to moderate or absent support for the corresponding split in bootstrap consensus trees depending on alignment (Fig. 1 and Table 2). Anc-SH illustrates the complementary pattern: the deep split is topologically stable, yet residue identities are more diffuse (lower mean PP) despite a confident overall fold (high pLDDT). For this system, therefore, structural plausibility does not imply a uniquely resolved evolutionary context. Conversely, robust topology does not guarantee high residue-level certainty at deep nodes. The composite BS–PP–pLDDT index used here is best viewed as a compact decision-support summary of these three dimensions, while still requiring inspection of each component for alignment-sensitive nodes such as Anc-HeR (Fig. 5). Consistent with this interpretation, increasing UFBoot2 replicates and varying random seeds yielded stable support for Anc-SzR and Anc-SH, whereas Anc-HeR remained sensitive and could alternate between recovery and collapse in the consensus depending on alignment and resampling settings (Table S4). Our multimeric predictions further support the biological plausibility of the reconstructed sequences.

AlphaFold-Multimer favored a trimeric assembly for Anc-SzR and a dimeric assembly for Anc-HeR, mirroring the known oligomeric tendencies of extant SzRs and HeRs (*20* –*25*) (Fig. 6 and Table S10). Notably, the preferred oligomeric states were supported by high interface scores (ipTM) only for the corresponding stoichiometries, while alternative assemblies scored substantially lower. In this case study, these predictions strengthen the interpretation that indel-aware full-length ancestors preserve not only monomeric 7TM folds but also EM elements that are compatible with oligomerization interfaces.

A primary contribution of this work is that the computational reconstructions translate into experimentally tractable proteins. Anc-SzR and Anc-HeR were expressed essentially “as reconstructed”, using only minimal His_6_ tags from standard pET vectors, without solubility fusions, truncations, or stabilizing point mutations. Both constructs yielded visibly pigmented *E. coli* pellets under induction with all-*trans* retinal (Fig. 7A), and purified proteins showed rhodopsin-like absorption maxima consistent with their clades (Fig. 7B). This closes a practical loop that is rarely demonstrated for full-length 7TM ASR: in this SzR–HeR case study, sequence-level support (PP), structure-level confidence (pLDDT), and biochemical tractability (retinal binding and visible spectra) converge on the same reconstructed ancestors.

Placing this result in context, binary gap–coding strategies similar in spirit have been applied previously to microbial rhodopsins to probe ancient photic niches; however, prior work primarily emphasized TM-core inferences (e.g., spectral tuning) and did not evaluate full-length extra-membrane architecture or experimental tractability via protein resurrection (*48*). Our results extend these computational reconstructions by showing that full-length, indel-aware ancestors can be expressed as retinal-binding holoproteins, and Anc-SzR additionally showed CCCP-sensitive light-driven alkalinization in a whole-cell pH assay, whereas Anc-HeR showed no detectable net pumping signal under the same conditions, consistent with the prevailing view that extant HeRs are not canonical light-driven ion pumps (Fig. 8).

Finally, we briefly note that Anc-HeR is sensitive not only to alignment choices but also to pipeline/model combinations, which helps interpret the supporting analyses in Table S3. In our comparisons, IQ-TREE-based reconstructions using Q.pfam+R7 (and robustness checks using LG+C60+F+R8) produced Anc-HeR sequences that were mutually similar across alignments. In contrast, a classical RAxML–PAML reconstruc-tion under an LG-based model generated an Anc-HeR sequence that was far more divergent (Table S3). In this system, the discrepancy is plausibly driven primarily by model differences (profile-based versus LG-type) as much as by software per se; the key point for the case study is that the HeR ancestor is a “fragile” node whose inferred sequence can shift substantially under reasonable alternative modelling choices, while Anc-SzR is comparatively robust. Practically, this motivates treating Anc-HeR as conditionally reliable and explicitly reporting sensitivity analyses for such nodes, rather than assuming that a single pipeline provides a definitive answer.

Several limitations delimit the scope of this case study. First, AlphaFold confidence metrics are strong indicators of structural plausibility but do not replace experimental validation of fold, oligomeric state, and function. Second, our HeR and SzR sampling differs in provenance and annotation depth, and some apparent EM diversity may reflect sampling biases. Third, the composite reliability index inherits assumptions from bootstrap resampling, ASR model specification, and structure prediction, and should be interpreted as an integrative summary rather than a standalone truth metric. Despite these caveats, the SzR–HeR system demonstrates a clear outcome: full-length, indel-aware ancestral reconstruction can yield compact 7TM sequences with coherent EM architecture, and at least two such ancestors can be expressed and recovered as retinal-binding rhodopsins without stabilizing engineering. These results establish a practical path for extending experimental ASR beyond trimmed TM cores and toward direct tests of how extra-membrane architecture co-evolves with the 7TM fold.

## Conclusions

In this study, we combined an indel-aware ASR workflow with AlphaFold-based confidence mapping to reconstruct full-length ancestral 7TM microbial rhodopsins, explicitly including not only the transmembrane helices but also extra-membrane regions. Crucially, the resulting ancestors were experimentally tractable: Anc-SzR and Anc-HeR were expressed essentially “as reconstructed” in *E. coli* and recovered as stable, retinal-binding holoproteins with rhodopsin-like visible absorption maxima. In addition, Anc-SzR showed CCCP-sensitive light-driven alkalinization in a whole-cell pH assay, whereas Anc-HeR showed no detectable net pumping signal under the same conditions, consistent with the prevailing view that extant HeRs are not canonical ion pumps. Together, these results show that, in the SzR–HeR system, full-length, indel-aware ASR can yield compact 7TM ancestors supported by both statistical (PP) and structural (pLDDT) evidence and can be taken forward for experimental testing by protein resurrection.

In microbial rhodopsins, functional diversification is often dominated by local changes around the retinal-binding pocket within the TM core, and extracellular loops may appear secondary at first glance. Nevertheless, our reconstructions suggest that substantial portions of EM segments can be inferred with only modestly reduced confidence relative to TM regions and recovered as structurally coherent motifs. These EM elements provide a consistent structural context for interpreting ancestral photochemistry, oligomerization, and potential protein–protein interactions, arguing against routine aggressive trimming in 7TM ASR. More broadly, many eukaryotic 7TM receptors, including GPCRs, rely on extracellular and cytoplasmic loops and C-terminal tails as primary determinants of ligand recognition and G-protein/arrestin coupling.

While extending the present approach to such systems will require careful attention to alignment depth, topology, and lineage-specific indel regimes, our results motivate gap-aware full-length reconstruction strategies in 7TM families where extra-membrane architecture is integral to function. One practical contribution of this work is to provide an explicitly documented, scriptable pipeline (ConsistASR: Consistency-aware ASR), including full control files for PAML-based reconstructions, so that each step from alignment to indel-aware ASR and structural evaluation can be replicated or adapted in other systems. We anticipate that the ConsistASR workflow introduced here will serve as a reproducible starting point for reconstructing and engineering ancestral 7TM proteins, treating both the transmembrane core and extra-membrane segments in a physically and statistically consistent manner.

## Accession Codes

UniProt accession codes are A0ACD6BAL3 for SzR4 and A0A2R4S913 for HeR 48C12. The accession codes for the proteins used in the phylogenetic analysis are summarized in the Supplementary Data.

## Supporting information

Additional experimental details and methods, including figures and tables.

## Acknowledgements

The authors would like to thank Taito Urui and Yu Iritani for helpful discussions and technical assistance. This research was supported by a Grants-in-Aid for Scientific Research to Y.M. (No. JP23H00285). The authors used a generative AI assistant (ChatGPT, GPT-5.1 Thinking; OpenAI) to assist with scriptwriting and to edit parts of the manuscript. (e.g. language polishing and restructuring of paragraphs). All scientific ideas, analyses, and conclusions are the authors’ own, and all AI-assisted text was carefully checked and, where necessary, revised by the authors.

## Supporting information

The following files are available free of charge.

- Supporting Information: Additional experimental details and methods, including figures of additional phylogenetic trees and AlphaFold predicted structures, and tables of sensitivity analysis and region-wise parameters (PDF).
- Supplementary Data: The sequence, alignment, phylogenetic, ancestral sequence, and structural data underlying the analyses in this manuscript, which are available at Zenodo (DOI: doi.org/10.5281/zenodo.19910435).

ConsistASR v1.0.1 is available on GitHub (github.com/harutois/ConsistASR) and archived at Zenodo (DOI: doi.org/10.5281/zenodo.18080328).

---

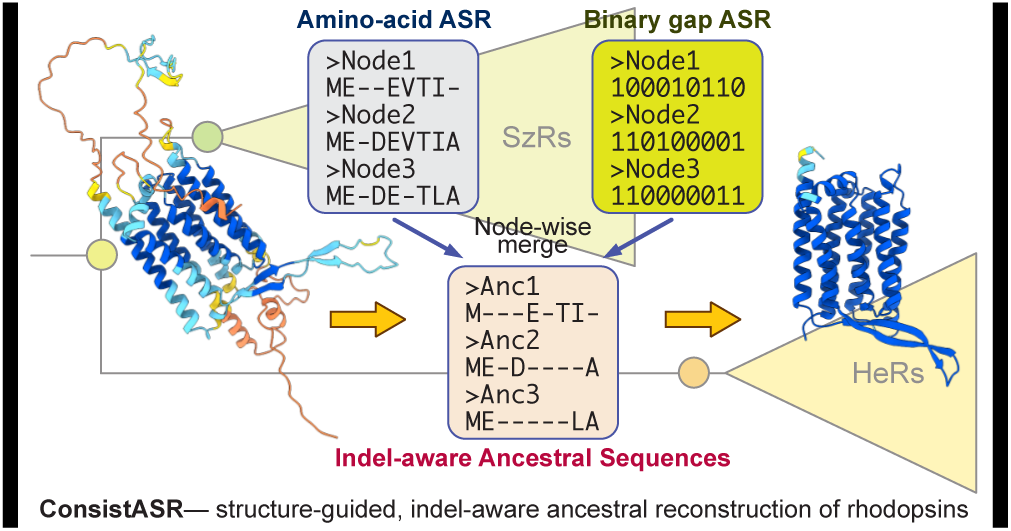
Table of Contents (TOC)/Abstract Graphic.

