## Additional experimental details and methods, including figures and tables. for "Resurrecting Full-length Ancestral Schizorhodopsins and Heliorhodopsins with Structure-guided, Indel-aware Sequence Reconstruction"

#### Supplementary Methods

##### S1. Detailed sequence retrieval and curation

###### S1.1 Database searches and seed sets

Previously characterized heliorhodopsins (HeRs) and schizorhodopsins (SzRs) were taken from Inoue et al. and Bulzu et al. (see main text). These seed sequences were used to initiate additional database searches:

- NCBI non-redundant protein database (nr) searched by BLASTP.
- MGnify clustered protein databases (version: 2024_04) searched by HMM-based profiles.

For MGnify, profile-based searches were performed using HMMER or the MGnify web interface; hit lists were filtered for sequences annotated as rhodopsins or containing seven predicted transmembrane helices.

###### S1.2 Length and motif filters

Length and mandatory-residue filters were implemented as a simple shell pipeline using `seqkit` and `awk`. As a generic first pass, we removed obviously short sequences from the combined HeR+SzR candidate set (`HeR_SzR_raw.fasta`):

```
# Example: length filter (here MINLEN = 150 aa)
seqkit fx2tab --name --length HeR_SzR_raw.fasta \
  | awk -v minlen=150 '$2 >= minlen { print $1 }' \
  > HeR_SzR_ids_len150.txt

seqkit grep -f HeR_SzR_ids_len150.txt HeR_SzR_raw.fasta \
  > HeR_SzR_len150.fasta
```

For the final curated sets, length thresholds were applied separately per family, using the generic script with family-specific settings (`MINLEN = 200` for HeRs, `MINLEN = 150` for SzRs).

An optional "mandatory residue" filter was implemented as a simple pattern search in `seqkit`, mirroring the `AA_FILTER` option in the generic `filter_cluster_generic.sh` script. For example, to keep only sequences containing at least one lysine in a given family-specific window around the Schiff-base site, we used:

```
# Example residue filter: keep sequences containing at least one K
seqkit grep -s -r -p "[K]" HeR_SzR_len150.fasta \
  > HeR_SzR_len150_KR.fasta
```

In practice, the precise positions corresponding to the canonical Schiff-base lysine in TM7 were determined from multiple sequence alignments, and the presence of the Schiff-base lysine at the aligned TM7 position was explicitly verified for the final curated sets. The shell commands above illustrate the underlying scripting logic; the complete, parameterized implementation is provided in the `filter_cluster_generic.sh` script in the ConsistASR repository.

##### S1.3 Redundancy reduction and diversity pruning

In the finalized dataset used in the main analyses, redundancy reduction was applied at 95% pairwise identity with CD-HIT, retaining one representative per cluster. The multi-step commands below illustrate an exploratory workflow for progressively collapsing near-identical sequences; the exact command history and thresholds used for the published dataset are recorded in the ConsistASR repository.

```
# Multi-step CD-HIT clustering of the length- and motif-filtered set
cd-hit -i HeR_SzR_len150_KR.fasta -o HeR_SzR_c99.fasta \
       -c 0.99 -n 5 -d 0 -T 8 -M 0

cd-hit -i HeR_SzR_c99.fasta -o HeR_SzR_c97.fasta \
       -c 0.97 -aS 0.8 -g 1 -d 0 -T 8 -M 0

cd-hit -i HeR_SzR_c97.fasta -o HeR_SzR_c95.fasta \
       -c 0.95 -aS 0.8 -g 1 -d 0 -T 8 -M 0
```

Tree-based diversity pruning was then applied to the 95% clustered set, following the same steps hard-coded in `filter_cluster_generic.sh`. Briefly, we aligned the 95% set and inferred a fast approximate tree:

```
# Alignment and preliminary tree from the 95% set
mafft --auto HeR_SzR_c95.fasta > HeR_SzR_c95.aln.fasta

FastTree -wag -gamma HeR_SzR_c95.aln.fasta \
  > HeR_SzR_c95.nwk
```

Treemmer v0.3 was used to prune near-identical tips while preserving deep branches until the target number of sequences was reached:

```
# Treemmer pruning to N = 225 sequences
python Treemmer.py HeR_SzR_c95.nwk -X 225 -np

# Treemmer writes a list of kept sequence IDs, e.g.:
# HeR_SzR_c95.nwk_trimmed_list_X_225

seqkit grep -f HeR_SzR_c95.nwk_trimmed_list_X_225 \
  HeR_SzR_c95.aln.fasta \
  > HeR_SzR_225.aln.fasta
```

The corresponding unaligned FASTA file used in the main analyses is `HeR_SzR_OG.fasta` in Supplementary data. The exact command sequence are provided in the ConsistASR repository; the examples above summarize the essential logic of the redundancy-reduction and diversity-pruning workflow.

#### S2. Multiple sequence alignment: implementation details

##### S2.1 Structure-aware alignments with PSI/TM-Coffee

Structure-aware multiple sequence alignments were generated with PSI/TM-Coffee in transmembrane mode on the T-Coffee server. In all cases, the input was the curated non-redundant set `HeR_SzR_OG.fasta` (Supplementary data), and UniRef100 was used as the homologous sequence database. Unless otherwise specified, all gap penalties and advanced options were left at their defaults.

Representative settings were:

- Input: `HeR_SzR_OG.fasta`
- Mode: "PSI/TM-Coffee" (transmembrane protein mode)
- Database: UniRef100 (TM-specific subset)
- All other parameters: server defaults (gap penalties, iteration scheme)

The alignment corresponding to this run is distributed as `HeR_SzR_OG.PSITM.fasta` in Supplementary data.

##### S2.2 MAFFT L-INS-i and FFT-NS-1

MAFFT v7.511 (online server) was used with the following strategies:

**L-INS-i:** L-INS-i alignments were generated on the MAFFT web server by selecting the "L-INS-i" strategy with default server settings. For reference, L-INS-i corresponds conceptually to local-pair iterative refinement (e.g. `-localpair` with extensive iterative refinement).

**FFT-NS-1:** FFT-NS-1 alignments were also generated on the MAFFT web server by selecting the "FFT-NS-1" strategy with default server settings.

The alignments corresponding to this run are distributed as `HeR_SzR_OG.LINSI.fasta` and `HeR_SzR_OG.FFTNS1.fasta` in Supplementary data. Alignments for HeR+outgroup and SzR+outgroup subsets were generated with the same strategies from the corresponding reduced FASTA files.

##### S2.3 trimAl-based trimming

Trimmed alignments were generated using trimAl v1.5 with the `gappyout` option:

```
trimal -in HeR_SzR_OG.PSITM.fasta \
       -out HeR_SzR_OG.PSITM.gappyout.fasta \
       -fasta \
       -gappyout
```

Trimmed versions were used for sensitivity analyses of model selection, tree topologies, and a conventional column-trimming baseline (see main text).

##### S2.4 Column-trimmed baseline alignments for ASR (trimAl gappyout)

To provide a conventional baseline for comparison with indel-aware reconstruction, we performed gap-unaware ASR on column-trimmed MSAs generated by trimAl `gappyout`. For each trimmed MSA, we re-estimated an ML tree under the best-fit model (typically Q.pfam+R7) and performed IQ-TREE marginal ASR on the resulting topology, analogously to the untrimmed analyses.

```
# Example (PSI/TM-Coffee gappyout baseline)
iqtree3 -s HeR_SzR_OG.PSITM.gappyout.fasta \
        -m Q.pfam+R7 \
        -bb 1000 -alrt 1000 \
        -nt AUTO \
        -o OG_WP_285271495,OG_WP_136361479,OG_WP_010903286 \
        -pre HeR_SzR_OG_QPFAMR7_PSITM_gappyout

iqtree3 -s HeR_SzR_OG.PSITM.gappyout.fasta \
        -m Q.pfam+R7 \
        -te HeR_SzR_OG_QPFAMR7_PSITM_gappyout.treefile \
        --ancestral \
        -nt AUTO \
        -o OG_WP_285271495,OG_WP_136361479,OG_WP_010903286 \
        -seed 12345 \
        -pre ASR_HeR_SzR_OG_QPFAMR7_PSITM_gappyout
```

Summary statistics for the gappyout baseline (length, mean PP, mean pLDDT) are reported in Table 3, and representative AlphaFold models are shown in Fig. S3.

#### S3. Model selection and tree inference in IQ-TREE

##### S3.1 ModelFinder runs

For each alignment, ModelFinder was called via IQ-TREE v3.0.1:

```
iqtree3 -s HeR_SzR_OG.PSITM.fasta \
        -m MFP \
        -bb 1000 \
        -alrt 1000 \
        -nt AUTO \
        -o OG_WP_285271495,OG_WP_136361479,OG_WP_010903286
```

The resulting `.log` file contains the BIC-based model ranking; for the main pipeline, Q.pfam+R7 (or closely related Q.pfam+R variants) was selected.

##### S3.2 ML tree searches under Q.pfam+R

Once the best-fit Q.pfam+R model was identified, ML tree searches were re-run with the model fixed, e.g.:

```
iqtree3 -s HeR_SzR_OG.PSITM.fasta \
        -m Q.pfam+R7 \
        -bb 1000 \
        -alrt 1000 \
        -nt AUTO \
        -o OG_WP_285271495,OG_WP_136361479,OG_WP_010903286 \
        -pre HeR_SzR_OG_QPFAMR7_PSITM
```

The output included:

- `.treefile`: ML tree with SH-aLRT and UFBoot2 support.
- `.contree`: UFBoot2 consensus tree.
- `.iqtree`: model-fit statistics, including BIC and log-likelihood.

##### S3.3 LG+C60+F+R8 mixture models

Profile-mixture trees were inferred under LG+C60+F+R8 on a Linux workstation:

```
iqtree3 -s HeR_SzR_OG.PSITM.fasta \
        -m LG+C60+F+R8 \
        -bb 1000 \
        -nt 10 \
        -o OG_WP_285271495,OG_WP_136361479,OG_WP_010903286 \
        -pre guide
```

```
iqtree3 -s HeR_SzR_OG.PSITM.fasta \
        -ft guide.treefile \
        -m LG+C60+F+R8+PMSF \
        -bb 1000 \
        -alrt 1000 \
        --tbe \
        -nt 10 \
        -o OG_WP_285271495,OG_WP_136361479,OG_WP_010903286 \
        -pre PMSF_C60
```

These runs were used only for robustness checks; the Q.pfam+R7 trees were used for the main ASR pipeline.

##### S3.4 UFBoot2 convergence checks

To assess the stability of ultrafast bootstrap support values for focal splits, UFBoot2 analyses were repeated with increasing numbers of replicates (IQ-TREE `-bb 1000`, `-bb 2000`, and `-bb 5000`). For MAFFT L-INS-i at 5,000 replicates, repeated the analysis with an alternative random seed (12345 and 67890) to assess stochastic variability. The resulting consensus support values are summarized in Table S4.

#### S4. IQ-TREE ancestral sequence reconstruction

##### S4.1 ASR command templates

Ancestral reconstructions were carried out with IQ-TREE using ML topologies inferred under the corresponding Q.pfam+R model:

```
iqtree3 -s HeR_SzR_OG.PSITM.fasta \
        -m Q.pfam+R7 \
        -te HeR_SzR_OG_QPFAMR7_PSITM.treefile \
        --ancestral \
        -o OG_WP_285271495,OG_WP_136361479,OG_WP_010903286 \
        -seed 12345 \
        -pre ASR_HeR_SzR_OG_QPFAMR7_PSITM
```

For each run, IQ-TREE produced:

- `.state` file: per-node, per-site marginal posterior probabilities.

Equivalent commands were used for MAFFT L-INS-i alignments with appropriate `-s`, `-te` and `-pre` arguments.

##### S4.2 Extant sequence reconstruction (ESR) proxy test

ESR proxy tests were carried out by replacing a target tip by a proxy subtree with two dummy leaves (`_A`, `_B`) and performing marginal ASR at the proxy internal node under the baseline setting (PSI/TM-Coffee; Q.pfam+R7). Proxy MSAs and trees were generated with `make_esr_proxy.py`. Dummy branch length was set to 500, where ESR identity scores were observed to plateau in exploratory tests (10–2000). Reconstruction accuracy was evaluated using `score_esr.py` and `score_esr_by_region.py` (Table 5, S9).

```
# Example (SzR4)
python make_esr_proxy.py \
  --msa HeR_SzR_OG.PSITM.fasta \
  --tree HeR_SzR_OG_QPFAMR7_PSITM.treefile \
  --target SzR_AM_5_00977 \
  --dummy-bl 500 \
  --out-prefix ESR_SzR4
```

```
iqtree3 -s ESR_SzR4.fasta \
        -m Q.pfam+R7 \
        -te ESR_SzR4.tree \
        --ancestral \
        -nt AUTO \
        -seed 12345 \
        --prefix ESR_SzR4_QPFAMR7
```

```
python score_esr_by_region.py \
  --orig-msa HeR_SzR_OG.PSITM.fasta \
  --target SzR_AM_5_00977 \
  --family SzR \
  --state ESR_SzR4_QPFAMR7.state
```

#### S5. Binary gap model and indel-aware masking

##### S5.1 One-command wrapper for indel-aware ASR

To make the indel-aware refinement reproducible and easy to apply, we provide a single wrapper script (run_indel_aware_iqtree.sh) in the ConsistASR package. This script takes as input

(i) the amino-acid MSA used for IQ-TREE ASR, (ii) the corresponding IQ-TREE ML topology (.treefile), and (iii) the IQ-TREE ancestral state file (.state),

and performs all steps needed to generate indel-aware ancestral FASTA files (with and without gaps) on the same topology.

A representative command for the PSI/TM-Coffee alignment was:

```
bash run_indel_aware_iqtree.sh \
  --msa   HeR_SzR_OG.PSITM.fasta \
  --tree  ASR_HeR_SzR_OG_QPFAMR7_PSITM.treefile \
  --state ASR_HeR_SzR_OG_QPFAMR7_PSITM.state \
  --prefix HeR_SzR_OG_QPFAMR7_PSITM \
  --outgroup "OG_WP_285271495,OG_WP_136361479,OG_WP_010903286"
```

This produces two key outputs:

- HeR_SzR_OG_QPFAMR7_PSITM_indel_withgap.fasta: indel-aware ancestor sequences retaining alignment gaps.
- HeR_SzR_OG_QPFAMR7_PSITM_indel_nogap.fasta: gap-stripped versions used for AlphaFold and summary statistics.

All intermediate files (binary alignment, RAxML-NG and RAxML-HPC logs and trees, node-mapping tables) are moved into a dedicated working directory (<prefix>_indel_work) and preserved for inspection.

Equivalent commands were run for the MAFFT L-INS-i alignment, changing only the -msa, -tree, -state and -prefix arguments to match the corresponding alignment and IQ-TREE output files.

##### S5.2 Internal steps performed by the wrapper

Internally, `run_indel_aware_iqtree.sh` executes the following steps, which are also available as standalone Python utilities in the ConsistASR package.

**(1) Recoding MSAs to binary gap matrices.** The amino-acid MSA is recoded to a binary (0/1) alignment in PHYLIP format using `msa_to_binary.py`; residues become 1 and gaps 0:

- Input: full-length amino-acid MSA (FASTA).
- Output: binary 0/1 MSA in strict PHYLIP format.

This preserves the taxon- and site-specific pattern of gaps while discarding amino-acid identity.

**(2) Evaluating the IQ-TREE topology under BIN+G.** The binary alignment is analyzed with RAxML-NG under a binary + gamma model (`BIN+G`) on the fixed IQ-TREE topology. This step re-optimizes branch lengths for the binary gap process while keeping the topology identical to the amino-acid tree. The best-scoring binary tree is written as `<prefix>_indel_eval.raxml.bestTree`.

**(3) Binary ancestral gap reconstruction with RAxML-HPC.** Using the optimized binary tree and alignment, RAxML-HPC is run in ancestral-reconstruction mode under a two-state model with among-site rate heterogeneity:

- Model: `-m BINGAMMA` (two-state + $\Gamma$).
- Topology: fixed to the RAxML-NG `bestTree` from step (2).
- Output: `RAxML_marginalAncestralStates.<prefix>` and `RAxML_nodeLabelledRootedTree.<prefix>`.

The ancestral-states file provides per-node, per-site binary calls (0/1) for all internal nodes on the IQ-TREE topology.

**(4) Node mapping between RAxML and IQ-TREE.** Because IQ-TREE and RAxML use different internal node labels, bipartition-based mapping is performed using `map_raxml_to_iqtree_nodes.py`. This utility reads both trees, identifies internal nodes by their bipartitions, and writes a mapping table linking IQ-TREE node IDs (including Anc-SzR, Anc-HeR and Anc-SH) to RAxML node IDs and their binary ancestral states.

**(5) Construction of indel-corrected ancestors.** Finally, `state_and_indel_to_fasta.py` merges IQ-TREE amino-acid ASR output (`.state`) with the mapped binary indel states:

- For each internal node and alignment column: if the binary state = 0 (gap), the site is removed for that ancestor.

- If the binary state = 1 (residue), the maximum-posterior amino acid and its PP from IQ-TREE are retained.

Concatenating retained sites in alignment order yields two FASTA outputs per run: a with-gap version (alignment coordinates preserved) and a gap-stripped version (used for AlphaFold predictions and PP/pLDDT summaries). All indel-aware ancestors analyzed in the main text are derived in this way.

Full source code of `run_indel_aware_iqtree.sh` and the underlying Python utilities (`msa_to_binary.py`, `map_raxml_to_iqtree_nodes.py`, `state_and_indel_to_fasta.py`) is included in the ConsistASR repository distributed with the Supplementary data.

##### S5.3 Indel-aware refinement for the classical RAxML–PAML pipeline

For the classical comparison pipeline (RAxML tree + PAML ASR; see Section "Classical RAxML+PAML workflow for comparison" in the main Methods), we applied the same two-stage indel-aware refinement as for the IQ-TREE-based workflow, but using PAML `.rst` files instead of IQ-TREE `.state` files.

A wrapper script (`run_indel_aware_paml.sh`) provided in the ConsistASR package automates the following steps:

1. recode the amino-acid MSA to a binary gap/non-gap matrix (1 = residue, 0 = gap);
2. evaluate the supplied RAxML topology on the binary alignment with RAxML-NG under a BIN+G model and write a best-scoring tree;
3. perform binary ancestral reconstruction with RAxML-HPC on the optimized binary tree;
4. map RAxML internal node IDs to PAML node IDs using the `.rst` tree; and
5. merge PAML amino-acid states and binary indel states to obtain gap-aware ancestral sequences (with-gap and gap-stripped) for all internal nodes.

A typical command was:

```
bash run_indel_aware_paml.sh \
  --msa    HeR_SzR_OG.FFTNS1.fasta \
  --tree   HeR_SzR_OG_FFTNS1.raxml.bestTree.tre \
  --rst    HeR_SzR_OG_FFTNS1.rst \
  --prefix HeR_SzR_OG_LGFG4_FFTNS1 \
  --outgroup "OG_WP_285271495,OG_WP_136361479,OG_WP_010903286"
```

Internally, the script calls `msa_to_binary.py` to generate the binary alignment, RAxML-NG and RAxML-HPC for binary ASR, and `map_raxml_to_paml_nodes_from_rst.py` plus `paml_state_and_indel_to_fasta.py` to produce PAML-based indel-aware ancestral FASTA files. The resulting “with-gap” and gap-stripped sequences were used for the RAxML–PAML comparison summarized in Table S3.

#### S6. AlphaFold predictions and data handling

##### S6.1 Single-chain models

Indel-uncorrected and indel-corrected sequences were submitted to an AlphaFold3 web server for single-chain structure prediction by providing one sequence as a single-chain input. For each submission, five models were generated, and only the top-ranked model was used for figures and summary statistics.

##### S6.2 Oligomeric models

For oligomeric-state analyses, indel-corrected sequences of Anc-SzR, Anc-HeR and Anc-SH were submitted as multi-sequence inputs to the same AlphaFold3-based web workflow to obtain homomeric assemblies (stoichiometries 1, 2, 3 and 5). In this web workflow, complex prediction is specified by providing multiple sequences in a single submission rather than by an explicit "Multimer" option.

For each stoichiometry, ipTM and pTM scores were extracted from the output (e.g. JSON or logs) and summarized in Table S10. Only the top-ranked model per stoichiometry was used in the figures.

#### S7. Mapping ASR confidence metrics onto AlphaFold models

##### S7.1 One-command wrapper for PP / pLDDT embedding

To visualize ASR posterior probabilities (PP) and to derive combined PP–pLDDT metrics on AlphaFold models, we used a wrapper script (`run_confmap_iqtree.sh`) included in the ConsistASR package.

Given (i) an IQ-TREE `.state` file, (ii) an indel-aware alignment containing the focal ancestor (with gaps retained), and (iii) the AlphaFold model directory for that ancestor, the script:

1. identifies the top-ranked AlphaFold model for the node from `*summary_confidences_*.json`,
2. converts the corresponding mmCIF file to PDB (preserving pLDDT in the B-factor field),
3. extracts per-site maximum posterior probabilities from the `.state` file for the specified node,
4. aligns these PP values to the indel-aware ancestral sequence, and
5. writes new PDB files in which B-factors encode PP, PP−pLDDT (scaled) or PP×pLDDT.

A typical usage within the directory containing the AlphaFold output for a given node (e.g. Anc-SzR) was:

```
bash run_confmap_iqtree.sh \
  --state  ASR_HeR_SzR_OG_QPFAMR7_PSITM.state \
  --node  Node3 \
  --withgap  HeR_SzR_OG_QPFAMR7_PSITM_indel_withgap.fasta \
  --outdir  confmap
```

Here, `-state` points to the IQ-TREE `.state` file used for ASR, `-node` specifies the focal internal node label, and `-withgap` refers to the indel-aware alignment FASTA in which that node appears as a gapped sequence (the `_indel_withgap.fasta` output from the indel-aware pipeline). The `-outdir` argument controls where the derived PDB files and summary statistics are written.

##### S7.2 Internal processing steps and outputs

Internally, `run_confmap_iqtree.sh` performs the following operations, using small Python utilities distributed with ConsistASR.

**(1) Selection of the AlphaFold model.** The script scans for `*summary_confidences_*.json` files in the current directory, optionally preferring those whose filenames contain the node name. Among these candidates, it selects the model with the highest `ranking_score` (or, if unavailable, the highest mean pLDDT) and infers the corresponding mmCIF file (`<prefix>_model_<index>.cif`).

**(2) CIF→PDB conversion with pLDDT in B-factors.** The selected mmCIF file is converted to PDB using an available converter (`phenix.cif_as_pdb`, `gemmi` or `pdb_fromcif`), ensuring that AlphaFold pLDDT values are preserved in the B-factor field. This PDB (e.g. `Node3_plddt_bfactor.pdb`) serves as the baseline structure in which B-factors represent pLDDT.

**(3) Extraction of per-site PP values.** Site-wise PP values for the specified node are extracted directly from the IQ-TREE `.state` file by scanning rows corresponding to that node and taking, at each alignment column, the maximum posterior over the 20 amino acids. These values are written as a simple text file (one PP per line), which forms the residue-level PP profile used in subsequent mapping and statistics.

**(4) Extraction of the indel-aware aligned sequence.** The ancestral sequence for the same node is retrieved from the indel-aware alignment (with gaps) using its FASTA header. This FASTA entry preserves the alignment positions and gap pattern after binary indel masking and therefore matches the coordinate system used for the PP vector.

**(5) Embedding PP into B-factors.** A helper script (`map_confidence_to_bfactor.py`) reads the baseline AlphaFold mmCIF, the aligned ancestral FASTA, and the PP values, and produces a new PDB in which the B-factor of each C$\alpha$ atom encodes PP on a 0–100 scale. This PDB (e.g. `Node3_pp_bfactor.pdb`) is used for visualizing residue-wise ASR confidence.

**(6) Derivation of PP−pLDDT and PP×pLDDT maps.** Using the PP-mapped and pLDDT-mapped PDBs, the script computes for each C$\alpha$ atom:

- **PP−pLDDT**: the difference between PP and pLDDT, scaled from $[-100, 100]$ to $[0, 100]$ for convenient colouring;
- **PP×pLDDT**: the product of PP and pLDDT (both on 0–100 scales), rescaled to [0,100].

Two additional PDB files are written in which B-factors encode these quantities (e.g. `Node3_ppminusplddt_bfactor.pdb` and `Node3_ppxplddt_bfactor.pdb`). These were used to generate the color-mapped surfaces shown in Fig. 4C and S4.

**(7) C$\alpha$-based summary statistics.** Finally, the script computes simple C$\alpha$-only statistics (mean, minimum, maximum and median) for each metric across the ancestor:

- site-wise PP (from the `.state` file),
- pLDDT (from the baseline PDB),
- PP (as mapped to B-factors),
- PP$-$pLDDT (scaled), and
- PP$\times$pLDDT.

These values are written to a log file (e.g. `Node3_conf_CA_stats.log`) and form the basis for the node-level summary metrics (mean PP, mean pLDDT, mean PP$\times$pLDDT, PP$-$pLDDT) reported in the main text and in Supplementary tables.

Full source code for `run_confmap_iqtree.sh` and the associated Python utilities is provided in the ConsistASR repository distributed with the Supplementary data.

##### S7.3 Application to PAML-based ancestors

For PAML-based ancestral reconstructions in the classical RAxML–PAML pipeline, we used an analogous wrapper (`run_confmap_paml.sh`) to embed site-wise posterior probabilities from PAML `.rst` files into AlphaFold models.

Conceptually, the workflow mirrors the IQ-TREE-based `run_confmap_iqtree.sh` procedure (Section S7.1): the script (i) selects the top-ranked AlphaFold model from `*summary_confidences_*.json`, (ii) converts the mmCIF file to PDB with pLDDT in B-factors, (iii) extracts per-site PAML posterior probabilities for a specified internal node from the `.rst` file, (iv) aligns these PP values to the corresponding indel-aware ancestral sequence in the PAML `withgap` FASTA, and (v) produces PDB files in which B-factors encode PP, scaled PP$-$pLDDT, or PP$\times$ pLDDT, together with C$\alpha$-only summary statistics.

A typical command in the directory containing the AlphaFold output for a given PAML node (e.g. 233) was:

```
bash run_confmap_paml.sh \
  --rst  HeR_SzR_OG_LGFG4_FFTNS1.rst \
  --node  233 \
  --withgap  HeR_SzR_OG_LGFG4_FFTNS1_indel_withgap.fasta \
  --outdir  confmap
```

Here, `-node` specifies the integer PAML node index as used in the `.rst` file; the script internally converts this to a label (`Node233`) and searches for the corresponding AlphaFold model. The resulting PP-, PP$-$pLDDT- and PP$\times$ pLDDT-mapped PDB files and log of C$\alpha$ statistics are written to the directory specified by `-outdir`. Full source code for `run_confmap_paml.sh` and its helper scripts (`extract_pp_from_paml_rst.py`, `map_confidence_to_bfactor.py`) is included in the ConsistASR repository.

#### S8. OPM-based TM/EM mapping

OPM entries corresponding to SzR4 (PDB 7e4g) and HeR 48C12 (PDB 6uh3) were downloaded, and residue-wise membrane annotations were mapped onto Anc-SzR and Anc-HeR via the PSI/TM-Coffee or MAFFT L-INS-i alignments. A higher-resolution structure of the same HeR (PDB 6su3) was used only for illustrative structural overlays; TM/EM assignments and region-level statistics are based on the OPM entry for PDB 6uh3.

#### S9. Classical RAxML+PAML workflow

##### S9.1 RAxML-NG tree inference

For the classical comparison, ML trees were inferred with RAxML-NG under LG+G4:

```
raxml-ng --all \
         --msa HeR_SzR_OG.FFTNS1.fasta \
         --model LG+F+G \
         --bs-metric tbe \
         --tree rand{1} \
         --bs-trees 1000 \
         --outgroup OG_WP_285271495,OG_WP_136361479,OG_WP_010903286 \
         --seed 12345 \
         --prefix HeR_SzR_OG_FFTNS1_LGFG4 \
```

Best ML trees (`*.raxml.bestTree`) were then used as fixed topologies for PAML.

##### S9.2 PAML codeml settings

PAML v4.10.7 codeml was run with a control file similar to:

```
seqfile = HeR_SzR_OG.FFTNS1.phy
treefile = HeR_SzR_OG_FFTNS1_LGFG4.raxml.bestTree
outfile = codeml_HeR_SzR_OG_FFTNS1.out

noisy = 3
verbose = 1
runmode = 0

seqtype = 2      * amino acid sequences
aaRatefile = lg.dat

model = 2       * Empirical + F
```

```
Mgene = 0

fix_alpha = 0
alpha = 0.5
ncatG = 4

fix_kappa = 1
kappa = 2
fix_omega = 1
omega = 1

cleandata = 0
RateAncestor = 1
```

The `rst` output file was parsed to extract ancestral sequences at nodes corresponding to Anc-SzR, Anc-HeR and Anc-SH and compared to IQ-TREE-based ancestors (Table S3).

#### S10. Detailed expression and purification conditions

##### S10.1 Culture conditions and induction

*E. coli* C43(DE3) cells harboring pET-21a(+)/Anc-SzR or pET-15b/Anc-HeR were grown on LB agar plates containing 50 $\mu$g/mL ampicillin. Single colonies were used to inoculate 4 mL LB pre-cultures, which were grown for 3 h at 37 °C with shaking (200 rpm). Pre-cultures were then used to inoculate 200 mL LB medium to an initial $OD_{600}$ of ∼0.05.

Cells were grown at 37 °C and 200 rpm until $OD_{600}$ reached ∼1.0, at which point protein expression was induced by addition of IPTG to a final concentration of 1 mM. All-*trans*-retinal was added simultaneously to a final concentration of 10 $\mu$M from a concentrated ethanol stock. For condition screens, cultures were incubated after induction under three conditions:

- 37 °C for 4 h
- 37 °C for 20 h
- 25 °C for 20 h

For preparative work, the 25 °C, 20 h condition was used, as it consistently yielded the strongest pigmentation.

##### S10.2 Membrane preparation

Cells were harvested by centrifugation at 6,000 $\times g$ for 10 min at 4 °C and resuspended in 50 mM Tris-HCl, 5 mM $MgCl_2$, pH 8.0. Cell suspensions were disrupted by sonication on ice (Taitec, VP-30S). Unbroken cells and debris were removed by low-speed centrifugation at 8,000 $\times g$ for 10 min at 4 °C. Membranes were collected by ultracentrifugation at 100,000 $\times g$ for 1 h at 4 °C (Hitachi, himac CS 150 GXII).

##### S10.3 Solubilization and affinity purification

Membrane pellets were resuspended in 50 mM MES-NaOH, 300 mM NaCl, 5 mM $MgCl_2$, 5 mM imidazole, pH 6.5, containing 1.5% (w/v) $n$-dodecyl-$\beta$-D-maltoside (DDM) and incubated at 4 °C for 1 h with gentle agitation. Insoluble material was removed by ultracentrifugation at 100,000 $\times g$ for 1 h at 4 °C. The supernatant containing solubilized rhodopsins was loaded onto a prepacked $Co^{2+}$-affinity column (Talon crude 5 mL, GE Healthcare) equilibrated in wash buffer (50 mM MES-NaOH, 300 mM NaCl, 50 mM imidazole, 5 mM $MgCl_2$, pH 6.5, 0.1% DDM).

After sample loading, the column was washed with at least five column volumes of wash buffer, and bound proteins were eluted with a linear imidazole gradient (50–500 mM) in elution buffer (50 mM Tris-HCl, 500 mM NaCl, 5 mM $MgCl_2$, pH 7.0, 0.1% DDM). Chromatography was performed on an ÄKTA pure system (GE Healthcare). Pigmented fractions were pooled and concentrated as needed, then buffer-exchanged into 50 mM Tris-HCl, 500 mM NaCl, pH 7.0, containing 0.05% DDM for spectroscopic measurements.

##### S10.4 UV–visible spectroscopy

UV–visible absorption spectra of purified, detergent-solubilized Anc-SzR and Anc-HeR were recorded at room temperature on a Shimadzu UV-3150 spectrophotometer using a 10 mm-path quartz cuvette. Spectra were scanned from 300 to 700 nm; the corresponding buffer was used for baseline subtraction.

#### S11. Whole-cell pH assay for light-driven proton transport

Light-driven ion-transport activity was assessed in whole cells by monitoring external pH changes upon illumination, following a commonly used assay for microbial rhodopsins. *E. coli* C43(DE3) cells expressing Anc-SzR or Anc-HeR were harvested by centrifugation, washed with 300 mM NaCl (no buffering agent), and resuspended in the same solution. Cell suspensions were adjusted to the same $OD_{600}$ before measurements. Aliquots (3.0 mL) of the cell suspension were transferred to a measurement cell equipped with a magnetic stir bar.

The suspension temperature was maintained at 25 °C using a circulating water bath (EYELA, UC-55N) while continuously stirring. External pH was monitored continuously with a pH meter (HORIBA, F-72) using an immersion-type pH electrode placed directly in the cell suspension. Baseline pH was recorded under dark conditions for 5 min, followed by illumination for 5 min at 540 nm, and then a post-illumination dark period for 10 min. For measurements shown as raw traces (Fig. S6), this dark–light–dark protocol was repeated for a second cycle on the same suspension (i.e., two consecutive cycles). Illumination at 540 nm was provided using a xenon lamp light source (Asahi Spectra Co., Ltd., MAX-303) equipped with a 540 nm bandpass filter. Illumination was performed using the same optical setup and settings across all measurements.

Where indicated, the protonophore carbonyl cyanide m-chlorophenyl hydrazone (CCCP) was added to the same suspension to a final concentration of 30 µM from a concentrated stock solution, mixed thoroughly, and incubated for 20 min in the dark prior to repeating the same illumination protocol. pH traces were recorded continuously throughout the experiment at a sampling interval of 2 s.

#### S12. Software environment

Unless otherwise stated, analyses were performed under macOS (64-bit) using bash and GNU core utilities. LG+C60 mixture-model calculations were run on a 64-bit Linux workstation due to their higher computational demands. Python scripts were written for Python 3 and rely on standard scientific libraries including `Biopython`. Exact software versions, command histories and configuration files are recorded in the ConsistASR repository and the accompanying Zenodo archive.

### Supporting Tables

**Table S1:** Additional dataset, alignment, and model combinations used for sensitivity analyses.

| Dataset | Alignment / trimming | $N$ sequences | Alignment length (residues) | Substitution model | Notes |
|---|---|---|---|---|---|
| HeR+SzR+outgroup | PSI/TM-Coffee / trimAl `gappyout` | 228 | 246 | Q.pfam+R7 | BIC-best model on moderately trimmed alignment (ModelFinder, IQ-TREE). |
| HeR+SzR+outgroup | MAFFT L-INS-i / trimAl `gappyout` | 228 | 245 | Q.pfam+R7 | Same best-fit family as untrimmed alignments; used to confirm robustness of model choice. |
| HeR+SzR+outgroup | PSI/TM-Coffee (untrimmed) | 228 | 449 | LG+C60+F+R8 (PMSF) | Profile-mixture model with improved BIC and log-likelihood relative to Q.pfam+R7; used only for sensitivity analyses. |
| HeR+SzR+outgroup | MAFFT L-INS-i (untrimmed) | 228 | 421 | LG+C60+F+R8 (PMSF) | Same profile-mixture model applied to the alternative alignment. |
| HeR+SzR+outgroup | MAFFT FFT-NS-1 (untrimmed) | 228 | 455 | LG+F+G4 | "Classical" ASR pipeline used for comparison (RAxML tree + PAML ASR). Model chosen to mimic previous ASR studies rather than as BIC-best. |

Unless otherwise noted in the "Notes" column, models were selected as BIC-best candidates by ModelFinder in IQ-TREE. LG+C60+F+R8 rows correspond to more complex profile-mixture models with better BIC than Q.pfam+R7, used here only for sensitivity analyses. The LG+F+G4 model in the MAFFT FFT-NS-1 / RAxML–PAML pipeline was chosen to emulate classical ASR workflows rather than by formal BIC comparison.

**Table S2:** Mean PP, mean pLDDT, and composite score for key ancestors under different pipelines.

| Ancestor | Alignment | Model / pipeline | Mean PP | Mean pLDDT | Composite score | Length (residues) |
|---|---|---|---|---|---|---|
| | PSI/TM-Coffee | Q.pfam+R7 / IQ-TREE | 81.6 | 95.5 | 77.3 | 206 |
| Anc-SzR | PSI/TM-Coffee | LG+C60+F+R8 / IQ-TREE | 80.3 | 95.3 | 75.0 | 206 |
| | FFT-NS-1 | LG+F+G4 / RAxML+PAML | 84.1 | 96.0 | NA[a] | 205 |
| | PSI/TM-Coffee | Q.pfam+R7 / IQ-TREE | 90.6 | 95.9 | 47.1 | 259 |
| Anc-HeR | PSI/TM-Coffee | LG+C60+F+R8 / IQ-TREE | 84.4 | 93.6 | 0.0[b] | 259 |
| | FFT-NS-1 | LG+F+G4 / RAxML+PAML | 88.0 | 94.3 | NA[a] | 263 |
| | PSI/TM-Coffee | Q.pfam+R7 / IQ-TREE | 54.9 | 93.2 | 51.6 | 206 |
| Anc-SH | PSI/TM-Coffee | LG+C60+F+R8 / IQ-TREE | 50.9 | 94.4 | 47.9 | 205 |
| | FFT-NS-1 | LG+F+G4 / RAxML+PAML | 56.1 | 94.2 | NA[a] | 205 |

Values are shown on a 0–100 scale. PP and pLDDT are averaged over C$\alpha$ atoms in indel-corrected ancestors. RAxML+PAML pipeline uses MAFFT FFT-NS-1, LG+F+G4, and indel-aware correction as described in the main text.

[a] Composite scores are defined only for IQ-TREE pipelines, where UFBoot2 support values are available for the focal split. For the RAxML+PAML pipeline, conventional nonparametric bootstrap values were not converted into composite scores because they are not directly comparable to UFBoot2 frequencies.

[b] Composite scores are defined as $100 \times b \times \overline{\mathrm{PP} \times \mathrm{pLDDT}}$, where $b_{\mathrm{UFBoot2}}$ is the UFBoot2 frequency of the focal split. When a branch is collapsed in the UFBoot2 consensus tree (i.e. the split is absent), $b_{\mathrm{UFBoot2}} = 0$ and the composite score is 0 by definition, even if $\overline{\mathrm{PP} \times \mathrm{pLDDT}}$ is high.

**Table S3:** Pairwise amino-acid identity (%) for Anc-HeR under different models and pipelines.

| Alignment<br>Model<br>Pipeline | PSI/TM-Coffee<br>Q.pfam+R7<br>IQ-TREE | MAFFT L-INS-i<br>Q.pfam+R7<br>IQ-TREE | PSI/TM-Coffee<br>LG+C60+F+R8<br>IQ-TREE | MAFFT FFT-NS-1<br>LG+F+G4<br>RAxML+PAML |
|---|---|---|---|---|
| PSI/TM-Coffee<br>Q.pfam+R7<br>IQ-TREE | 100 | 88.8 | 92.7 | 52.2 |
| MAFFT L-INS-i<br>Q.pfam+R7<br>IQ-TREE | 88.8 | 100 | 89.2 | 53.1 |
| PSI/TM-Coffee<br>LG+C60+F+R8<br>IQ-TREE | 92.7 | 89.2 | 100 | 49.8 |
| MAFFT FFT-NS-1<br>LG+F+G4<br>RAxML+PAML | 52.2 | 53.1 | 49.8 | 100 |

Comparisons reflect both model and software/pipeline differences; the RAxML+PAML reconstruction uses LG+F+G4, whereas IQ-TREE reconstructions use Q.pfam+R7 or LG+C60+F+R8.

**Table S4:** UFBoot2 convergence assessment for focal ancestral splits under different alignments.

| Alignment | UFBoot2 setting | Anc-SzR | Anc-HeR | Anc-SH |
|---|---|---|---|---|
| MAFFT L-INS-i | 1,000 (seed 12345) | 96 | 0[a] | 100 |
| | 2,000 (seed 12345) | 98 | 0[a] | 100 |
| | 5,000 (seed 12345) | 97 | 47 | 100 |
| | 5,000 (seed 67890) | 95 | 0[a] | 100 |
| PSI/TM-Coffee | 1,000 (seed 12345) | 97 | 61 | 100 |
| | 2,000 (seed 12345) | 97 | 64 | 100 |
| | 5,000 (seed 12345) | 97 | 62 | 100 |

Ultrafast bootstrap support values (UFBoot2) were computed from IQ-TREE consensus trees (`.contree`) using increasing numbers of bootstrap replicates (IQ-TREE: `-bb 1000-5000`) and, for MAFFT L-INS-i at 5,000 replicates, an alternative random seed. Values are reported for the three focal nodes (Anc-SzR, Anc-HeR, Anc-SH). When the focal split was not recovered in the UFBoot2 consensus (collapsed into a local polytomy), support is reported as 0 and annotated as "collapsed".

[a] The focal Anc-HeR split was absent from the UFBoot2 consensus tree (collapsed).

**Table S5:** Region-wise mean posterior probabilities (PP) for Anc-SzR and Anc-HeR (indel-corrected).

| Region | Anc-SzR (PSI/TM-Coffee) | Anc-SzR (MAFFT L-INS-i) | Anc-HeR (PSI/TM-Coffee) | Anc-HeR (MAFFT L-INS-i) |
|---|---|---|---|---|
| N-terminal EM | 68.7 | 88.0 | 70.1 | 70.7 |
| TM1 | 75.0 | 71.1 | 88.7 | 89.0 |
| EM1 (TM1–TM2) | 79.6 | 75.5 | 88.7 | 87.7 |
| TM2 | 81.3 | 78.3 | 95.1 | 96.6 |
| EM2 (TM2–TM3) | 79.8 | 79.1 | 88.1 | 92.2 |
| TM3 | 92.2 | 93.5 | 95.8 | 98.9 |
| EM3 (TM3–TM4) | 70.8 | 64.7 | 99.7 | 99.8 |
| TM4 | 85.3 | 85.7 | 92.0 | 93.7 |
| EM4 (TM4–TM5) | 86.3 | 82.3 | 98.8 | 97.9 |
| TM5 | 82.7 | 82.7 | 90.1 | 93.3 |
| EM5 (TM5–TM6) | 69.7 | 67.3 | 75.2 | 76.1 |
| TM6 | 92.2 | 88.1 | 92.7 | 89.9 |
| EM6 (TM6–TM7) | 92.0 | 86.1 | 96.2 | 96.4 |
| TM7 | 80.5 | 78.2 | 96.7 | 97.0 |
| C-terminal EM | 64.5 | 89.6 | 87.5 | 92.4 |

EM segments correspond to OPM-guided extra-membrane regions between TM helices (e.g. EM1 = TM1–TM2 segment).
Values are reported on a 0–100 scale.

**Table S6:** Region-wise mean pLDDT for Anc-SzR and Anc-HeR (indel-corrected).

| Region | Anc-SzR (PSI/TM-Coffee) | Anc-SzR (MAFFT L-INS-i) | Anc-HeR (PSI/TM-Coffee) | Anc-HeR (MAFFT L-INS-i) |
|---|---|---|---|---|
| N-terminal EM | 82.8 | 91.5 | 82.3 | 80.0 |
| TM1 | 97.6 | 95.3 | 97.7 | 97.5 |
| EM1 (TM1–TM2) | 94.1 | 71.0 | 97.0 | 96.1 |
| TM2 | 97.5 | 93.4 | 97.9 | 97.6 |
| EM2 (TM2–TM3) | 89.2 | 89.9 | 94.8 | 95.2 |
| TM3 | 96.5 | 96.1 | 97.9 | 96.4 |
| EM3 (TM3–TM4) | 94.9 | 94.9 | 97.4 | 95.2 |
| TM4 | 97.1 | 97.3 | 98.3 | 97.9 |
| EM4 (TM4–TM5) | 94.4 | 94.5 | 96.7 | 96.1 |
| TM5 | 97.8 | 97.8 | 97.7 | 95.0 |
| EM5 (TM5–TM6) | 88.6 | 88.4 | 89.5 | 73.0 |
| TM6 | 97.5 | 96.9 | 96.4 | 89.5 |
| EM6 (TM6–TM7) | 95.2 | 95.4 | 96.1 | 96.0 |
| TM7 | 97.8 | 96.8 | 97.6 | 94.7 |
| C-terminal EM | 86.4 | 81.8 | 92.8 | 88.9 |

EM segments correspond to OPM-guided extra-membrane regions between TM helices (e.g. EM1 = TM1–TM2 segment).
Values are reported on a 0–100 scale.

**Table S7:** Region-wise mean (PP × pLDDT) for Anc-SzR and Anc-HeR (indel-corrected).

| Region | Anc-SzR (PSI/TM-Coffee) | Anc-SzR (MAFFT L-INS-i) | Anc-HeR (PSI/TM-Coffee) | Anc-HeR (MAFFT L-INS-i) |
|---|---|---|---|---|
| N-terminal EM | 55.9 | 81.1 | 57.8 | 57.1 |
| TM1 | 73.2 | 67.8 | 86.7 | 86.8 |
| EM1 (TM1–TM2) | 74.7 | 53.8 | 86.0 | 84.2 |
| TM2 | 79.4 | 74.0 | 93.1 | 94.3 |
| EM2 (TM2–TM3) | 71.1 | 71.1 | 83.6 | 87.8 |
| TM3 | 88.9 | 89.7 | 93.8 | 95.3 |
| EM3 (TM3–TM4) | 67.1 | 61.4 | 97.0 | 95.0 |
| TM4 | 82.8 | 83.3 | 90.4 | 91.7 |
| EM4 (TM4–TM5) | 81.3 | 77.5 | 95.5 | 94.2 |
| TM5 | 80.9 | 80.9 | 88.2 | 88.7 |
| EM5 (TM5–TM6) | 62.8 | 60.0 | 67.3 | 55.7 |
| TM6 | 89.8 | 85.4 | 89.4 | 80.8 |
| EM6 (TM6–TM7) | 87.6 | 82.1 | 92.5 | 92.5 |
| TM7 | 78.8 | 75.9 | 94.3 | 91.8 |
| C-terminal EM | 58.2 | 73.4 | 81.6 | 82.1 |

EM segments correspond to OPM-guided extra-membrane regions between TM helices (e.g. EM1 = TM1–TM2 segment).
Values are reported on a 0–100 scale.

**Table S8:** Region-wise mean (PP − pLDDT) for Anc-SzR and Anc-HeR (indel-corrected).

| Region | Anc-SzR (PSI/TM-Coffee) | Anc-SzR (MAFFT L-INS-i) | Anc-HeR (PSI/TM-Coffee) | Anc-HeR (MAFFT L-INS-i) |
|---|---|---|---|---|
| N-terminal EM | 42.9 | 48.3 | 43.9 | 45.4 |
| TM1 | 38.7 | 37.9 | 45.5 | 45.8 |
| EM1 (TM1–TM2) | 42.8 | 52.3 | 45.8 | 45.8 |
| TM2 | 41.9 | 42.5 | 48.6 | 49.5 |
| EM2 (TM2–TM3) | 45.3 | 44.6 | 46.6 | 48.5 |
| TM3 | 47.9 | 48.7 | 48.9 | 51.3 |
| EM3 (TM3–TM4) | 38.0 | 34.9 | 51.2 | 52.3 |
| TM4 | 44.1 | 44.2 | 46.8 | 47.9 |
| EM4 (TM4–TM5) | 45.9 | 43.9 | 51.1 | 50.9 |
| TM5 | 42.4 | 42.4 | 46.2 | 49.2 |
| EM5 (TM5–TM6) | 40.6 | 39.5 | 42.9 | 51.5 |
| TM6 | 47.3 | 45.6 | 48.2 | 50.2 |
| EM6 (TM6–TM7) | 48.4 | 45.4 | 50.1 | 50.2 |
| TM7 | 41.4 | 40.7 | 49.5 | 51.2 |
| C-terminal EM | 39.1 | 53.9 | 47.4 | 51.8 |

EM segments correspond to OPM-guided extra-membrane regions between TM helices (e.g. EM1 = TM1–TM2 segment).
Values are reported on a 0–100 scale.

**Table S9:** Extant sequence reconstruction (ESR) accuracy across additional representative tips (global identity only). Dummy branch length was fixed to 500, where ESR estimates were saturated.

| Family | Target ID | Global identity (%) |
|---|---|---|
| SzR | SzR_Ga0105045_102227662 (AntR) | 68.4 |
| SzR | SzR_AM_5S_00009 (SzR1) | 85.1 |
| SzR | SzR_SAMEA_2622822 (SzR2) | 93.6 |
| SzR | SzR_TE_S2S_00499 (SzR3) | 98.6 |
| SzR | SzR_AM_5_00977 (SzR4) | 73.3 |
| HeR | HeR_AVZ43932 (HeR 48C12) | 76.6 |
| HeR | HeR_WP_100389406 | 92.9 |
| HeR | HeR_HWR63593 | 90.2 |
| HeR | HeR_ZSoct5m_G40_00154 | 85.7 |
| HeR | HeR_MEK9135620 | 80.3 |

**Table S10:** AlphaFold-Multimer predictions of oligomeric assemblies for ancestral rhodopsins.

| Alignment | Ancestor | Stoichiometry | ipTM | pTM |
|---|---|---|---|---|
| PSI/TM-Coffee | Anc-SzR | Monomer | – | 0.94 |
| | | Dimer | 0.23 | 0.56 |
| | | Trimer | 0.88 | 0.90 |
| | | Pentamer | 0.51 | 0.59 |
| | Anc-HeR | Monomer | – | 0.93 |
| | | Dimer | 0.81 | 0.86 |
| | | Trimer | 0.32 | 0.51 |
| | | Pentamer | 0.42 | 0.52 |
| | Anc-SH | Monomer | – | 0.92 |
| | | Dimer | 0.25 | 0.56 |
| | | Trimer | 0.67 | 0.73 |
| | | Pentamer | 0.26 | 0.38 |
| MAFFT L-INS-i | Anc-SzR | Monomer | – | 0.94 |
| | | Dimer | 0.23 | 0.56 |
| | | Trimer | 0.85 | 0.87 |
| | | Pentamer | 0.29 | 0.42 |
| | Anc-HeR | Monomer | – | 0.93 |
| | | Dimer | 0.79 | 0.84 |
| | | Trimer | 0.27 | 0.46 |
| | | Pentamer | 0.29 | 0.41 |
| | Anc-SH | Monomer | – | 0.93 |
| | | Dimer | 0.31 | 0.60 |
| | | Trimer | 0.77 | 0.81 |
| | | Pentamer | 0.16 | 0.30 |

For each ancestor, alignment, and stoichiometry, we report the predicted interface confidence (ipTM) and overall complex confidence (pTM). Scores are shown on a 0–1 scale; monomeric predictions report pTM only.

#### Supporting Figures

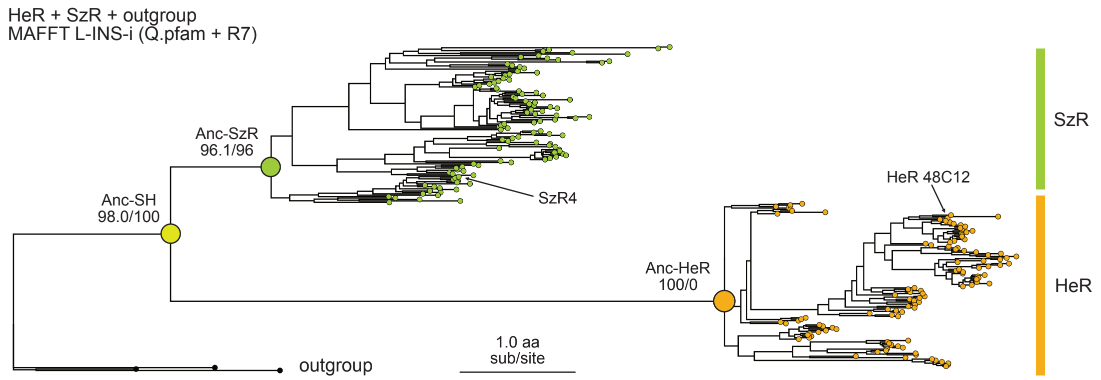


**Figure S1: Maximum-likelihood tree for HeR, SzR and outgroup rhodopsins based on the MAFFT L-INS-i alignment.** Full HeR+SzR+outgroup phylogeny inferred with IQ-TREE from the MAFFT L-INS-i alignment under the Q.pfam+R7 model (best-fit by BIC). The tree is rooted with three outgroup microbial rhodopsins, and the heliorhodopsin (HeR) and schizorhodopsin (SzR) clades are indicated. The positions of Anc-SzR, Anc-HeR and Anc-SH correspond to those highlighted in Fig. 1 of the main text. SH-aLRT and UFBoot2 support values at key internal branches are shown as reported by IQ-TREE. This figure provides the full-tree counterpart to the HeR-focused zoom in Fig. 1C, illustrating the overall stability of the SzR clade and the locally shortened branch leading to Anc-HeR under the MAFFT L-INS-i alignment.

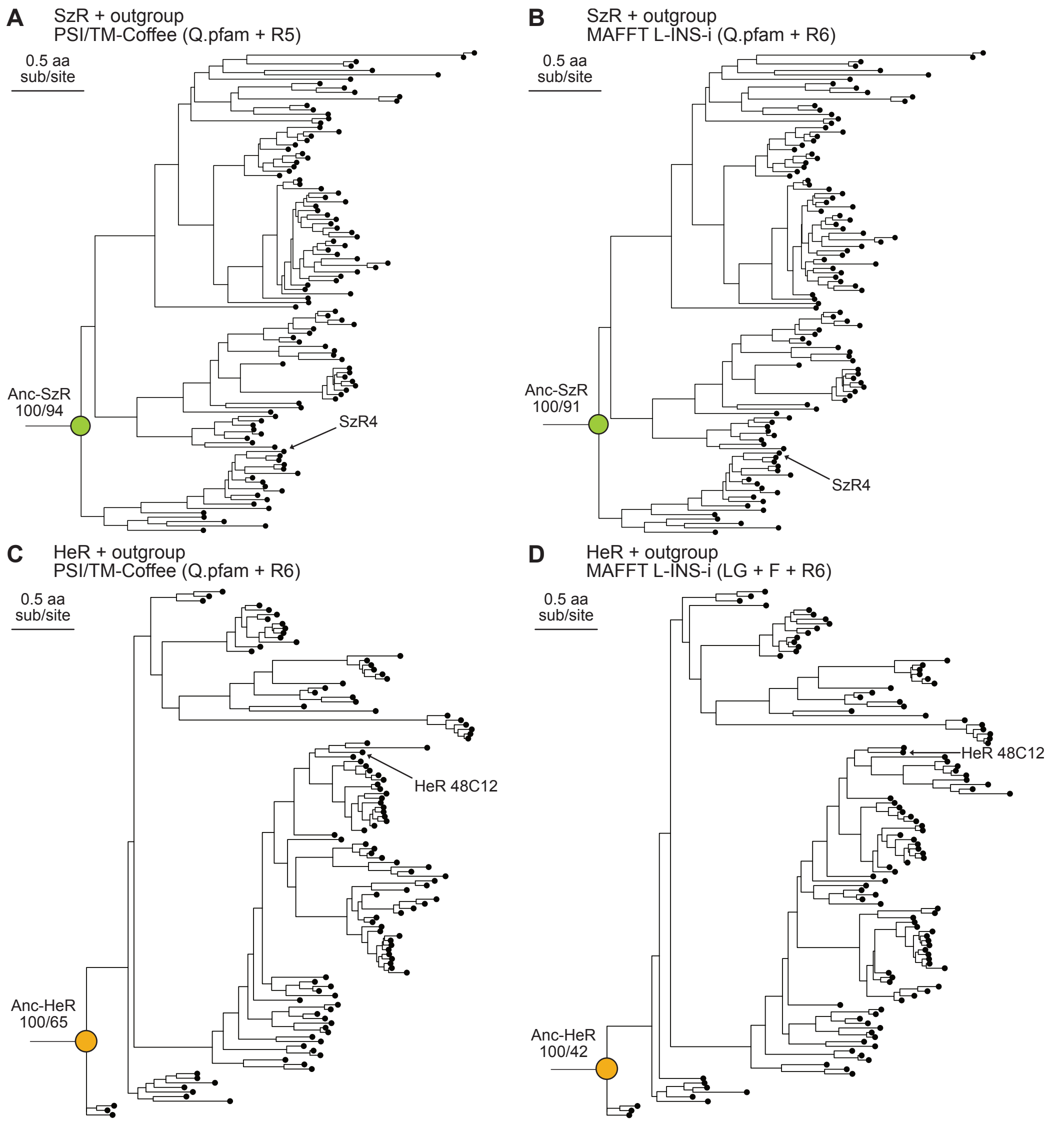


**Figure S2: Subset trees for SzR+outgroup and HeR+outgroup datasets under different alignment strategies.** Maximum-likelihood trees inferred with IQ-TREE for reduced datasets comprising either SzR+outgroup or HeR+outgroup sequences, using PSI/TM-Coffee (PSITM) or MAFFT L-INS-i (LINSI) alignments and the best-fit models selected by BIC. The ancestral nodes (Anc-SzR and Anc-HeR) are labeled together with their SH-aLRT/UFBoot2 support values. Extant structures used as templates (SzR4 and HeR 48C12) are also indicated. (A) SzR+outgroup tree based on the PSITM alignment. The Anc-SzR node is recovered with maximal SH-aLRT and UFBoot2, illustrating the robustness of the SzR clade to alignment strategy. (B) SzR+outgroup tree based on the LINSI alignment. Anc-SzR again shows very high support, confirming that the SzR ancestral split is stable across alternative alignments. (C) HeR+outgroup tree based on the PSITM alignment. Although the HeR clade is clearly separated from the outgroup, the internal branch leading to Anc-HeR is locally unstable and tends towards a near-polytomy in both the ML and consensus trees. (D) HeR+outgroup tree based on the LINSI alignment. Here, the best-fit model differs from the Q.pfam family and the Anc-HeR branch again shows reduced UFBoot2 support, highlighting that the HeR ancestral split is particularly sensitive to alignment details and model choice.

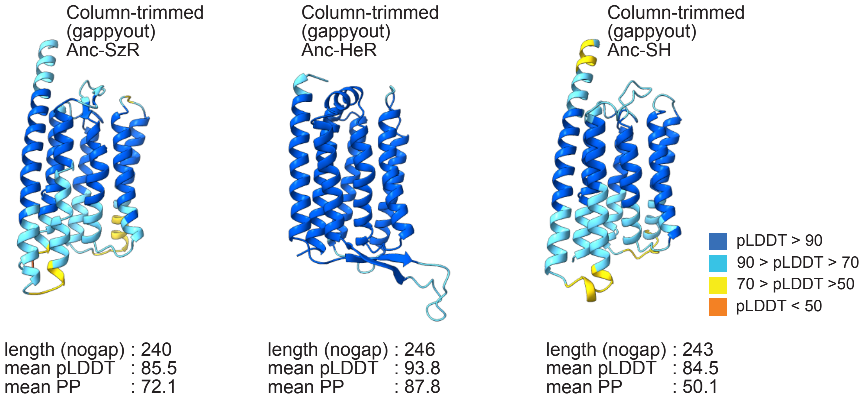


**Figure S3: AlphaFold models for the column-trimmed baseline (`gappyout`).** AlphaFold monomer models for ancestral sequences inferred from column-trimmed alignments generated with trimAl `gappyout` (no explicit indel treatment), shown for Anc-SzR, Anc-HeR, and Anc-SH under PSI/TM-Coffee. Structures are colored by per-residue pLDDT using the standard AlphaFold color scale. For each model, the gap-stripped sequence length (`nogap`), mean posterior probability (PP), and mean pLDDT are indicated (Cα-only). Compared with untrimmed gap-unaware ASR (Fig. 2A), column trimming reduces extreme overextension and improves overall confidence, but the resulting ancestors remain longer and/or less uniformly confident than the indel-aware, node-specific binary-masked ancestors (Fig. 2C and Table 3).

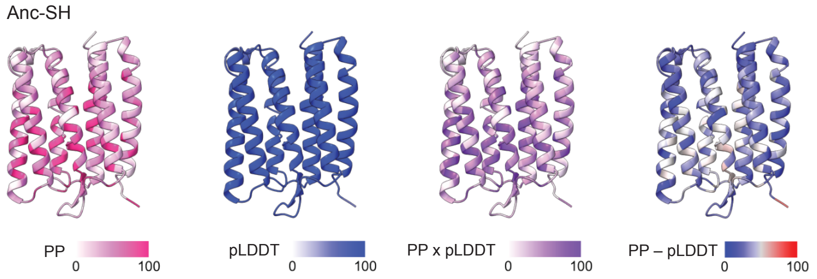


**Figure S4: Structure-based reliability maps for Anc-SH.** AlphaFold model of the indel-corrected Anc-SH sequence inferred from the PSI/TM-Coffee alignment, colored by four sequence–structure metrics: per-residue posterior probability (PP), per-residue pLDDT, the product PP × pLDDT, and the difference PP − pLDDT. PP and pLDDT values are mapped to a 0–100 scale and represented using the same color schemes as in Fig. 4C of the main text (red for high PP, blue for high pLDDT, purple for high PP × pLDDT, and a blue–grey–red scale for PP − pLDDT). Despite substantially lower mean PP than Anc-SzR and Anc-HeR, Anc-SH retains a well-defined 7TM fold with uniformly high pLDDT, illustrating that deep ancestral nodes can be structurally coherent yet residue-wise diffuse. This figure complements Fig. 4 by showing that the "topologically stable but sequence-ambiguous" character of Anc-SH is reflected directly in its site-wise PP and pLDDT profiles.

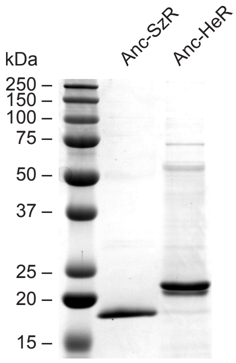


**Figure S5: SDS–PAGE analysis of detergent-solubilized Anc-SzR and Anc-HeR.** Purified proteins were separated on a 12% polyacrylamide gel and visualized by Coomassie staining. Lanes: molecular-weight ladder (kDa); lane 1, Anc-SzR; lane 2, Anc-HeR. Both ancestors migrate as essentially single bands with apparent molecular weights (∼18 and ∼22 kDa, respectively) lower than their calculated masses (24.7 and 31.5 kDa), an anomalous mobility commonly observed for multi-pass membrane proteins and not indicative of proteolysis. A faint higher-molecular-weight band is visible for Anc-HeR (lane 2), consistent with a minor oligomeric or detergent-associated species, and no prominent lower-molecular-weight fragments are detected.

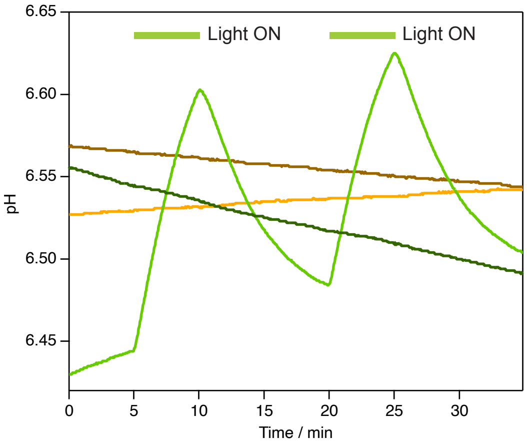


**Figure S6: Raw whole-cell pH traces underlying Fig. 8.** External pH was monitored for unbuffered *E. coli* suspensions (300 mM NaCl) expressing Anc-SzR or Anc-HeR during the same dark/light protocol as in Fig. 8, without baseline subtraction (raw pH). Two consecutive cycles are shown for each condition. Thick bars above the traces indicate illumination periods (540 nm). Traces are shown for Anc-SzR without CCCP (green) and with CCCP (30 µM; dark green), and for Anc-HeR without CCCP (orange) and with CCCP (dark orange).
